# *N*-Linked Glycosylation as a Driver of Tau Pathology and Neuronal Transmission

**DOI:** 10.64898/2025.12.01.690311

**Authors:** Tamara Basta, Morgane Herlory, Patrick Holland, Radmila Stojkovic, Wyatt Powell, Maciej A. Walczak

## Abstract

*N*-Linked glycosylation of Tau is implicated in Alzheimer’s disease, yet how it influences Tau propagation and neuronal uptake remains unclear. Here, we use chemically defined *N*-glycoforms of 2N4R Tau and the K18 repeat domain to map how glycan site and structure reprogram Tau-polyanion binding, seeding, and entry into human neurons. Site-specific GlcNAc or high-mannose glycans at Asn359 or Asn410 remodel heparin and RNA binding kinetics in biolayer interferometry assays, revealing avidity-dominated, slow-off heparin interactions that are dampened by *N*-glycans and faster RNA interactions that can be potentiated by small GlcNAc modifications. Cross-seeding experiments show that *N*-glycosylation selectively tunes nucleation compatibility between Tau proteoforms; GlcNAc at Asn410 in full-length Tau and small glycans at Asn359 in K18 most strongly accelerate aggregation. In HEK293T FRET biosensor cells and iPSC-derived cortical neurons, *N*-glycans enhance or suppress seeding and pHrodo-reported uptake in a site- and glycan-dependent manner, with Asn410 glycosylation markedly potentiating cellular seeding. Pharmacological inhibition with atropine, dynasore, and cytochalasin D further indicates that *N*-glycans rebalance muscarinic, dynamin-dependent, and actin-dependent uptake routes. Together, these data establish *N*-linked glycosylation as a tunable regulator of Tau trafficking and prion-like spread, coupling extracellular receptor usage to intracellular cofactor engagement, and reveals druggable nodes in uptake.

## INTRODUCTION

The progressive spread of misfolded Tau protein across anatomically connected brain regions is a defining feature of Alzheimer’s disease (AD) and related tauopathies, such as frontotemporal dementia and chronic traumatic encephalopathy.^1–3^ This pathological propagation is closely associated with cognitive decline and neuronal loss, yet the molecular mechanisms that govern Tau transmission remain incompletely understood. Tau is predominantly a cytosolic protein that stabilizes microtubules in neurons; however, in disease states, it undergoes a variety of PTMs, including phosphorylation, acetylation, ubiquitination, and glycosylation that can induce conformational changes and promote aggregation.^4^ While *O*-linked glycosylation of Tau has been documented for decades,^5–7^ evidence for *N*-linked glycosylation has only recently come to light.^8–12^ Early mass spectrometry studies of Tau from AD brains identified *N*-glycosylated species, particularly in the absence of signal peptides or transmembrane domains, suggesting an unconventional mechanism of Tau glycosylation or secretion.^13, 14^ Notably, aberrant *N*-glycosylation has been detected in hyperphosphorylated and aggregated Tau species, raising questions about its role in disease-associated conformations.^15^ More recent studies indicate that in apparently healthy controls, Tau contains triantennary *N*-linked glycans decorated with sialic acids at Asn359, whereas in AD samples, high-mannose glycans were observed at Asn410.^12^ One emerging hypothesis is that *N*-glycosylation facilitates misfolding, stabilization of toxic oligomers, and uptake by neighboring neurons via receptor-dependent mechanisms. Glycosylated Tau may exhibit altered interaction with surface receptors such as LRP1,^16^ heparan-sulfate proteoglycans (HSPGs),^17^ or carbohydrate-binding proteins (lectins),^18^ thereby modulating its endocytosis and intracellular trafficking. Additionally, glycosylated Tau may escape degradation^19^ or promote prion-like templating in recipient cells,^20^ contributing to the stereotyped progression of Tau pathology observed in Braak staging.

Despite these advances, the functional consequences of *N*-glycosylation in AD and on Tau biology remain poorly characterized.^21, 22^ Understanding how this modification influences Tau conformation, aggregation, secretion, and neuronal uptake is crucial for developing targeted strategies to block the spread of toxic Tau species.^23^ Because studying *N*-glycosylation resents formidable challenges in the control of glycan position and composition,^24^ we recently developed chemical and biochemical methods to selectively install *N*-linked glycans at the two confirmed positions in 2N4R Tau (Asn359 and Asn410) decorated with high-mannose structures.^25^ These studies revealed that in addition to changing in vitro biophysical properties of the protein such as the ability to undergo in vitro liquid-liquid phase separation, branched high-mannose glycans at the Asn410 position increase the propensity of Tau to undergo phosphorylation.^25, 26^ These results are consistent with the observations that the PHFs can be both hyperphosphorylated and *N*-glycosylated, and removal of *N*-linked glycans from PHFs by endoglycosidase F/N-glycosidase F results in reorganization into straight filaments.^10^ Based on these results, we wondered how glycosylation impacts the biophysical properties of Tau and how these changes translate into the cellular phenotypes, such as uptake and receptor involvement. Here, using in vitro aggregation assays and hiPSC-derived neurons, we map how discrete glycans reprogram Tau seeding efficiency and uptake mechanisms, revealing glycan-dependent switches that govern the efficiency and pathway of internalization.

## RESULTS

Prior studies indicate that the efficiency of heparin-induced seeding is diminished when large high-mannose glycans potentially block the growing fibrils or induce conformations that are not compatible with the unmodified protein.^25^ To determine whether *N*-glycosylation acts as a switch for Tau spread, we engineered site-specific glycoforms at Asn359 or Asn410 and compared their seeding potency (AF_t50_) and internalization pathways (Figures 1 and 2). This integrated in vitro-cellular approach allowed us to disentangle effects of glycan position, branching, and fibril state on nucleation kinetics. To this end, we first studied the efficiency of the glycosylated protein to propagate in an in vitro assay (Figure 2A). Pre-formed fibrils of K18, K18G, and K18M were used as seeds in a ThT heparin-induced fibrilization of K18 and K18M monomers. Cross-seeding analyses of K18 proteoforms bearing defined N359 glycans reveal a selective and directional network of interactions. or K18 monomers, K18C (GlcNAc_2_) seeds yielded the strongest acceleration (AF_t50_ median ≈ 5.2×, 4.7–5.6), outperforming homologous K18 seeds (≈ 3.0×, 2.8–3.2) and K18G seeds (≈ 3.7×, 3.3–4.1). In contrast, high-mannose K18M seeds produced little advancement in t₅₀ (≈ 1.0×, 0.9–1.0), despite a steep growth phase and high endpoints, indicating that bulky glycans on the seed can decouple amplitude from nucleation kinetics. For K18M monomers, the pattern becomes amplified: K18C, unmodified K18, and K18G seeds all drove very rapid nucleation with large acceleration factors (AF_t50_ medians ≈ 17.9×, 13.9×, and 13.2×, respectively), whereas high-mannose K18M seeds were substantially weaker (≈ 3.0×, 2.9–3.1). Together, these data show that small glycans at N359 on the seeds in K18 (GlcNAc/GlcNAc_2_) preserve or enhance the templating surface, supporting fast nucleation of both unmodified and high-mannose monomers, while bulky high-mannose on the seed impairs early nucleation, especially when the incoming monomer is also high-mannose. The forest plot on Figure 2C (median with 95% CI) visualizes this network and highlights an asymmetry: robust K18 → K18M but weak K18M → K18/K18M.

**Figure 1.**
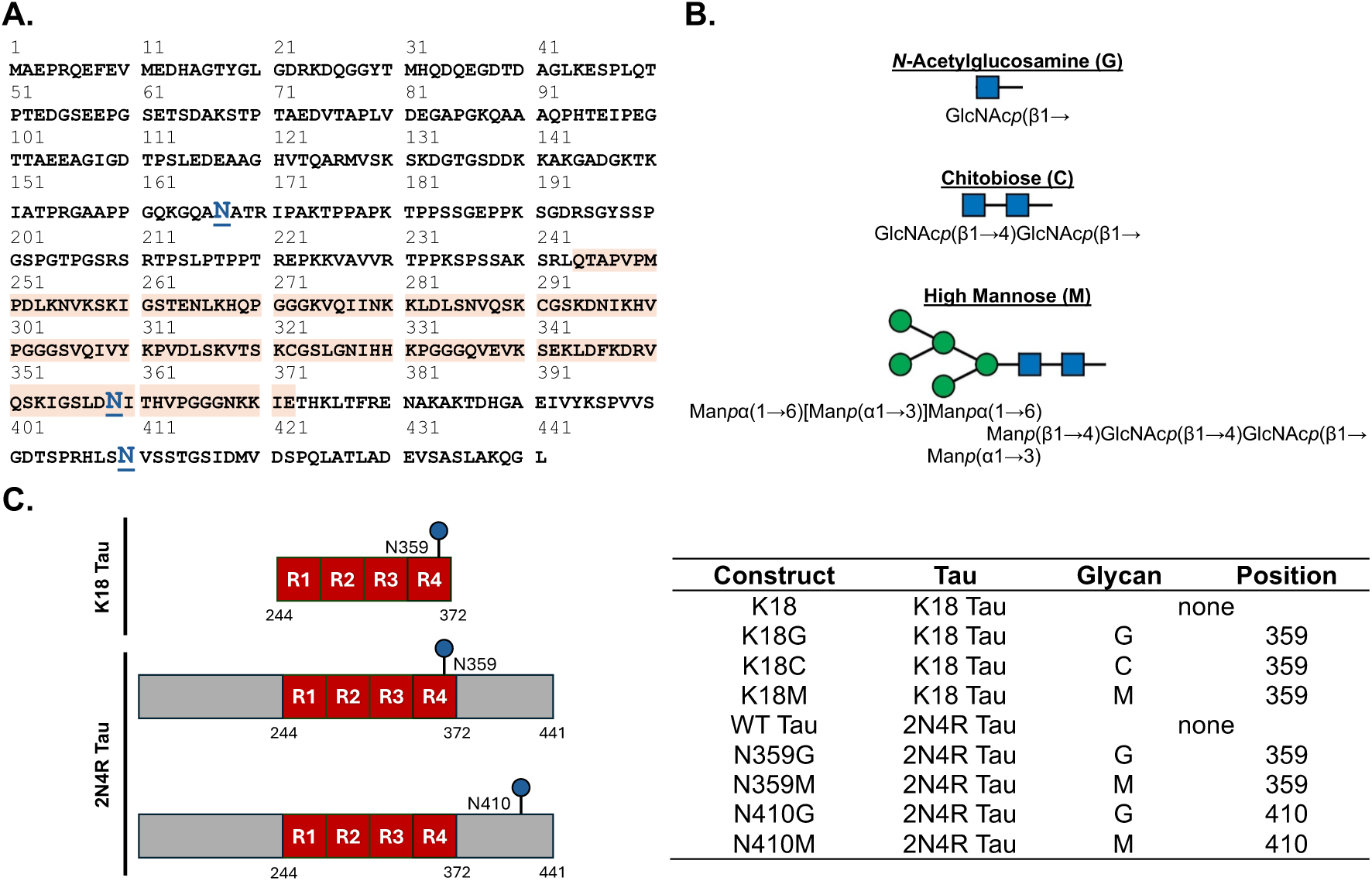
Overview of Tau glycoprotein constructs and their relevance to this study. **(A)** Primary amino acid sequence of 2N4R Tau with predicted (N167) and confirmed (N359 and N410) sites of *N*-linked glycosylation shown in blue. The red segment denotes the K18 Tau fragment. **(B)** Three classes of *N*-linked glycans attached at asparagine residues. *N* Acetylglucosamine (G) and chitobiose (C) serve as minimal model glycans that introduce limited structural perturbations to Tau, whereas the high-mannose heptasaccharide (M) with defined branching corresponds to the most abundant *N*-linked glycan characterized on PHFs from AD brain by Liu et al.^9^ **(C)** Summary of K18 and 2N4R Tau glycoproteins used in this study. Residue numbering refers to asparagine positions in full-length 2N4R Tau as defined in panel A.

**Figure 2.**
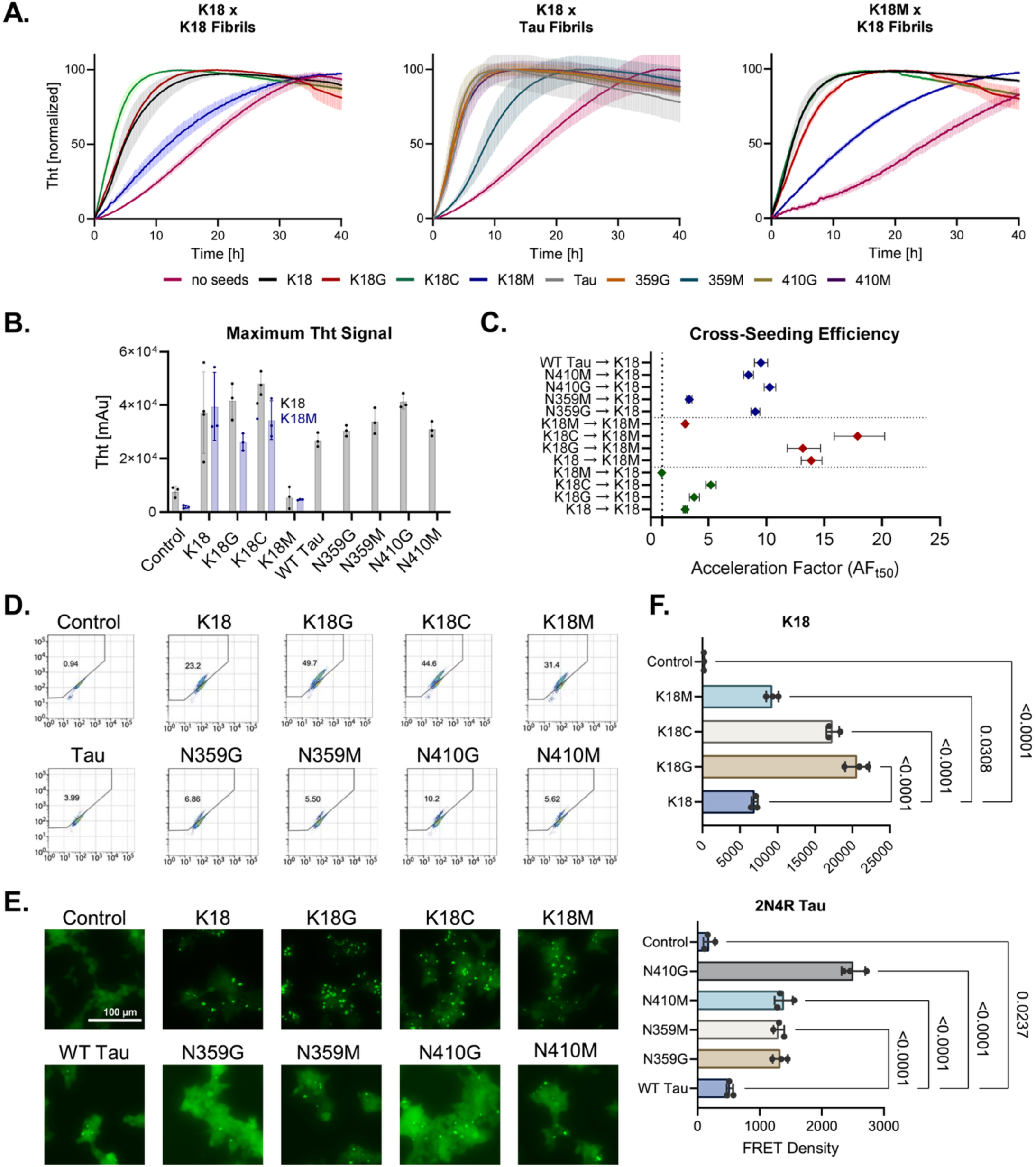
Seeding competency of Tau glycoprotein constructs in vitro and in cell-based biosensor assays. **(A)** ThT-monitored aggregation of K18 and K18M monomers seeded with pre-formed K18 or 2N4R Tau fibrils, quantified as relative fluorescence over time. **(B)** Maximum ThT fluorescence signal for each condition, quantified at the individual reaction maxima. **(C)** Cross-seeding efficacy expressed as an acceleration factor relative to unseeded reactions, where the acceleration factor corresponds to the ratio of t_50_ of unseeded versus seeded controls (t_50_,unseeded / t_50_,seeded). **(D)** FRET flow cytometry detection of seeding after transfection of K18 or 2N4R Tau proteoforms into HEK293T K18 CFP/YFP biosensor cells. (E) Representative confocal microscopy images of HEK293T CFP/YFP K18 biosensor cells transfected with vehicle, K18, or Tau proteoforms (20× objective; scale bar, 100 µm). **(F)** After seeding with vehicle, K18, or Tau proteoforms (100 nM, 24 h), integrated FRET density was quantified by flow cytometry. Vehicle denotes the lipofectamine-only negative control. Increases in FRET signal for glycosylated K18 and Tau proteoforms are shown relative to their respective WT constructs. n = 3 independent experiments for each proteoform; statistical significance was assessed by ordinary one-way ANOVA. **(G)** Representative flow cytometry scatter plots for each proteoform depicting FRET-positive cells (10,000 events analyzed per condition).

Next, we analyzed the aggregation efficacy of K18 in the presence of pre-formed 2N4R Tau seeds. In these assays, all full-length Tau seeds markedly accelerated K18 nucleation, but with a strong dependence on glycan identity and position. Tau N410G seeds were the most potent accelerators (AF_t50_ ≈ 10.3×), closely followed by GlcNAc at N359 (≈ 9.0×), unmodified WT Tau (≈ 9.5×), and high-mannose at N410 (≈ 8.4×). A Tau positive control produced the largest effect (≈ 11.6×). By contrast, high-mannose at N359 was the least effective full-length seed (≈ 3.3×), although it still accelerated K18 aggregation by ∼3× relative to unseeded reactions. Mechanistically, we propose that high-mannose at N359 sterically perturbs the docking surface or stabilizes a polymorph with reduced compatibility for early K18 monomer addition, whereas a small GlcNAc at N359 maintains a nucleation-competent interface. In contrast, both GlcNAc and high-mannose at N410 are largely tolerated, indicating that N410 resides outside the core templating surface recognized by K18 and can accommodate bulky glycans without major loss of cross-seeding efficiency.

To translate these observations into cellular context, we used HEK293T biosensor cell line expressing RD-CFP/YFP K18 (P301S) FRET pair as an indicator of intracellular aggregation.^27^ In the biosensor cells, we observed robust glycan-dependent enhancements in cellular seeding across both K18 and full-length 2N4R Tau at 50 nM aggregated seeds after 24 h (Figure 2D-2F). In the K18 panel, mono-GlcNAc at N359 produced the highest signal, followed by chitobiose K18C and high-mannose K18M, each significantly above unmodified K18 and vehicle yielding relative orders K18G > K18C > K18M > K18 ≫ control. In the 2N4R Tau panel, glycosylation at N410 yielded the strongest effects: N410-GlcNAc reached 2,508 A.U. (13.6× control), exceeding N410 high-mannose (1,388; 7.53×), N359-GlcNAc (1,331; 7.22×), and N359-high-mannose (1,307; 7.09×), all above unmodified WT Tau (2.81×) and vehicle. These effects align with (i) the sensitivity and specificity of the FRET-flow biosensor assay (femtomolar Tau seed detection) pioneered by Diamond and colleagues, validating that our integrated FRET density readout tracks intracellular templating by exogenous Tau seeds, and (ii) known mechanisms wherein HSPGs and co-receptors regulate aggregate uptake and seeding in this system.^27^ Mechanistically, there is a clear relationship between the site of glycosylation and templating efficiency: cryo-EM shows the AD PHF core spans residues 306–378, placing N359 within the β-core and N410 in the C-terminal “fuzzy coat,” consistent with our finding that N410-GlcNAc most strongly potentiates cellular seeding.^28^ Collectively, these data demonstrate that *N*-glycosylation enhances Tau seeding in cells in a site- and glycan-dependent manner, with N410-GlcNAc the standout modifier in full-length Tau and GlcNAc/GlcNAc_2_ most effective in K18, extending literature that links Tau glycosylation, HSPG/LRP1-mediated uptake, and prion-like propagation.^20^

Both, in vitro heparin-induced aggregation assays and the uptake of extracellular Tau are mediated by glycosaminoglycan, with heparin as the model polyanionic saccharide. To better understand how the polyanion-Tau interactions mediate the aggregation measured in Figure 2, we quantified how polyanion identity and site-specific *N*-glycosylation influence binding landscape across full-length 2N4R Tau and the repeat domain construct K18 (Figure 3). Heparin bound most tightly and with strikingly slow dissociation: 2N4R Tau-heparin exhibited K_D_ ≈ 14 nM with k_on_ ≈ 2.3×10^4^ M⁻¹s⁻¹ and k_off_ ≈ 3×10⁻⁴ s⁻¹ (residence time ∼55 min), whereas K18 bound more weakly (K_D_ ≈ 223 nM; residence time ∼4.2 min), consistent with avid, electrostatically driven multivalent contacts in the full-length protein. In 2N4R Tau, *N*-glycosylation at both asparagine positions (GlcNAc or high-mannose) uniformly reduced heparin affinity (K_D_ ≈ 37-79 nM), suggesting steric/electrostatic attenuation of avidity. In contrast, analogous modifications within K18 modestly tightened binding (K_D_ ≈ 133-176 nM vs 223 nM), implying local rearrangements in the repeat domain that stabilize productive contacts.

**Figure 3.**
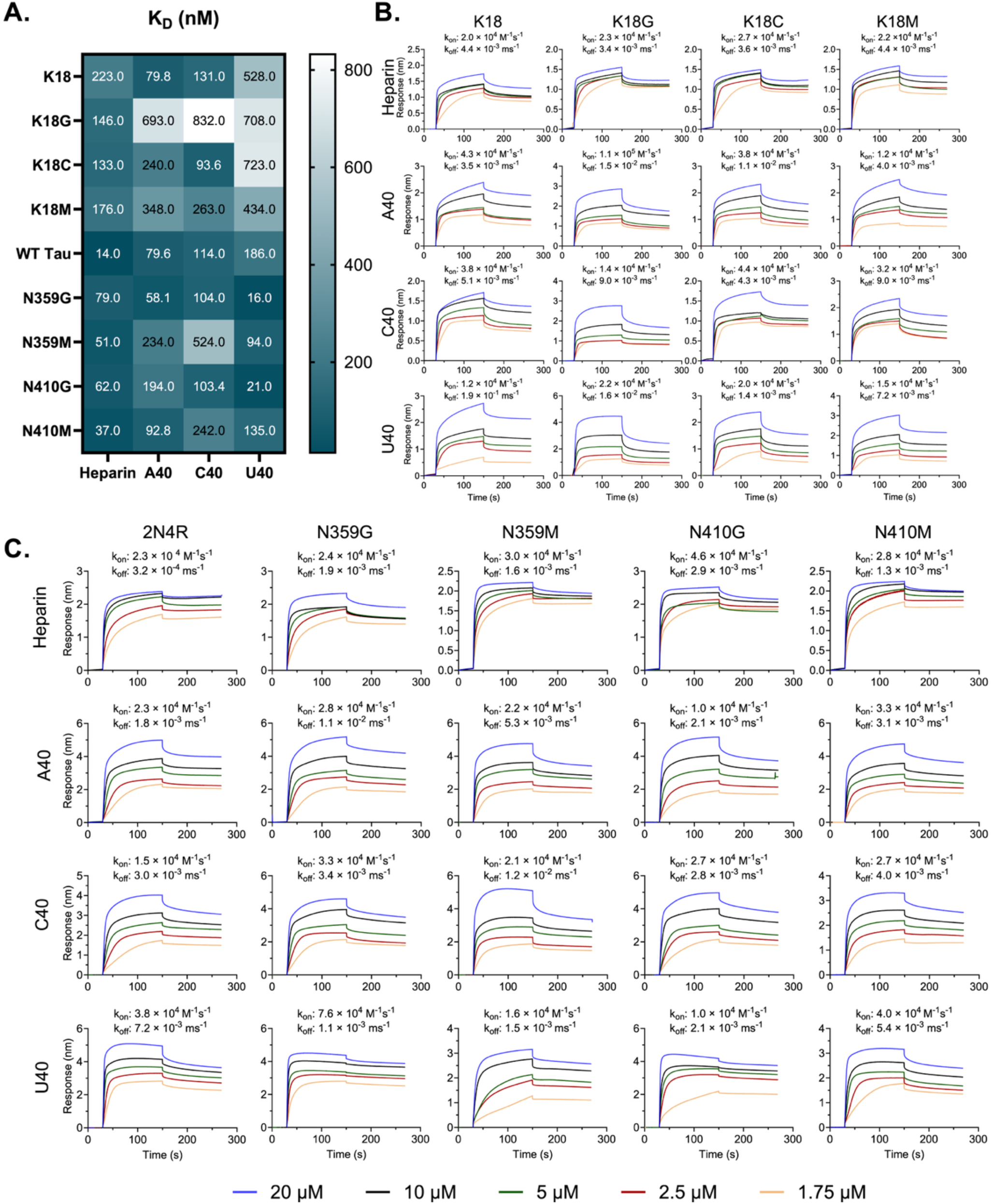
BLI analysis of K18 and Tau binding to heparin and RNA ligands. **(A)** Heat map of equilibrium dissociation constants (K_D_) obtained by BLI for K18 and Tau binding to heparin and RNA (A40, C40, U40). K_D_ values (nM) were derived from global fits across the full concentration series (20 µM–1.75 µM). **(B)** BLI binding curves for K18 proteoforms interacting with heparin and RNA. Experiments were performed in triplicate, and the shaded regions represent SD. After ligand immobilization, sensors were baselined for 30 s, followed by 120 s association and 120 s dissociation at each K18 concentration (20 µM–1.75 µM). **(C)** BLI binding curves for Tau proteoforms interacting with heparin and RNA ligands, measured under identical conditions as for K18. Kinetic parameters and K_D_ values reflect global fitting across all concentrations.

We next extended these studies to understand Tau-polyanion interactions with RNA. Lysine-rich microtubule-binding repeats (R1-R4) and adjacent basic regions drive electrostatic binding to polyanionic RNAs.^29–32^ Homopolymeric RNAs (A40, C40, U40) bound in the nanomolar-submicromolar range with broadly similar affinities but distinct sequence- and site-dependent modulation: for U40, GlcNAcylation of 2N4R Tau at Asn359 or Asn410 markedly strengthened binding (e.g., K_D_ ≈ 16-21 nM vs 186 nM for WT Tau), consistent with glycan-induced exposure or stabilization of RNA-interacting surfaces; high-mannose effects were smaller and mixed. Within A40 and C40 RNAs, variant preferences were evident (e.g., A40 favored N359G, K_D_ ≈ 58 nM; C40 favored K18C, K_D_ ≈ 94 nM). Together, these data reveal two mechanistic regimes: heparin-driven, slow-off avidity in full-length Tau that is dampened by bulky *N*-glycans, and faster, sequence-sensitive ssRNA interactions that can be potentiated by small, site-specific GlcNAcylation establishing a quantitative framework for how disease-relevant glycosylation and polyanion chemistry tune Tau binding kinetics. Contextually, our values align with and extend prior observations: SPR on repeat-domain fragments reported K18-heparin K_D_ ≈ 0.2 µM (vs K19 ≈ 70 µM) with binding strongly dependent on 6-O-sulfation and heparin chain length, whereas BLI with Tau immobilized has yielded apparent sub-nanomolar to picomolar affinities for heparin (e.g., ∼60 ± 30 pM), underscoring configuration- and avidity-driven tightening our 2N4R Tau-heparin K_D_ ≈ 14 nM thus falls within the expected low-nM/pM regime when multivalent contacts are accessible.^33, 34^ These trends dovetail with cell and biochemical studies showing that 6-O-sulfation is necessary and 3-O-sulfation enhances Tau-HS interactions and uptake, supporting our finding that bulky *N*-glycans dampen heparin binding by sterically and electrostatically attenuating access to clustered lysines that read out sulfate motifs. In contrast, 40-mer RNA homopolymers (A/C/U) bind full-length Tau with similar nanomolar avidity and a 2:1 model, yet only A40 efficiently drives strain formation matching our observation that small, site-specific GlcNAc at Asn359/Asn410 can potentiate RNA binding (notably to U40) without necessarily recapitulating heparin avidity, plausibly by shifting the conformational ensemble (e.g., relaxing proline-rich region (PRR)/microtubule-binding region (MTBR) contacts) to expose or geometrically align nucleic-acid interaction patches while avoiding the sulfate-pattern “key-lock” required for maximal HS avidity. Given that pathological *N*-glycosylation of Tau at Asn359/Asn410 has been detected in disease contexts, we speculate that these glycans could gate polyanion selectivity in vivo blunting HSPG-mediated uptake (reduced heparin avidity) while stabilizing intracellular RNA-Tau assemblies that influence seeding and strain maintenance, thereby coupling site-specific glycosylation to both extracellular trafficking and intracellular cofactor choice.

The interneuronal transmission of Tau is believed to contribute to the spread of the pathology across the brain.^35–37^ While the prior studies indicate the role of various receptors in assisting with this process, the role of post-translational modifications, including *N*-glycosylation is unknown. We used WTC11-derived i^3^Neurons, that were exposed to varying concentrations of monomeric and aggregated K18 and 2N4R Tau proteoforms labelled with a pH-sensitive dye, pHrodo red (Figures 3 and 4). We confirmed that each proteoform adjusted to 10% level of labeling show effectively identical fluorescence at pH 5.5, therefore allowing for a direct comparison of mean fluorescence intensities proportional to the proteoform concentration. Across WTC11-derived neurons, pHrodo-based time courses and end-point analyses showed that neuronal uptake/acidification is driven primarily by aggregation state and secondarily tuned by the N359 glycan, yielding a consistent rank order K18C > K18M > K18 > K18G for aggregated species at every dose tested (Figure 4). At 50 nM, aggregated proteoforms exhibited steep initial accumulation with mean early rates of 0.510 (K18C), 0.368 (K18M), 0.264 (K18G), and 0.150 RFU·h⁻¹ (K18) and convergent half-times near ∼12 h (K18C 11.75 h; K18G 12.17 h; K18 12.46 h; K18M 12.52 h), culminating in 24 h intensities of ∼98 (K18C), ∼96 (K18M), ∼69 (K18), and ∼54 RFU (K18G). Soluble counterparts remained near baseline (K18C ∼9.7, K18 ∼4.5, K18M ∼4.1, K18G ∼2.9 RFU). Early-window slopes (0-1 h and 1-3 h) and segmented AUCs (0-5 h vs 5-24 h) recapitulated these trends, indicating modest early capture but disproportionately larger late (5–24 h) contributions for aggregated species. Consistently, 24 h/3 h fold-changes at 50 nM were ∼10–15× for aggregated vs ∼2–4× for soluble. At 24 h, aggregated uptake exceeded soluble for every proteoform at 50, 150, and 250 nM, and large aggregation effects by standardized magnitude. Concentration-response at 24 h was approximately linear within 50-250 nM range for aggregates with smaller slopes for soluble species; preliminary E_max_/EC_50_ fits were consistent with no clear saturation in this range. These results establish that aggregation state is the dominant determinant of uptake/acidification, while glycan structure at N359 modulates the magnitude: chitobiose and high-mannose enhance aggregated uptake above non-glycosylated K18, whereas a single GlcNAc attenuates it. Controls (pHrodo-glycine and pHrodo-dextran conjugates) remained near baseline and replicate variability did not alter these rankings.

**Figure 4.**
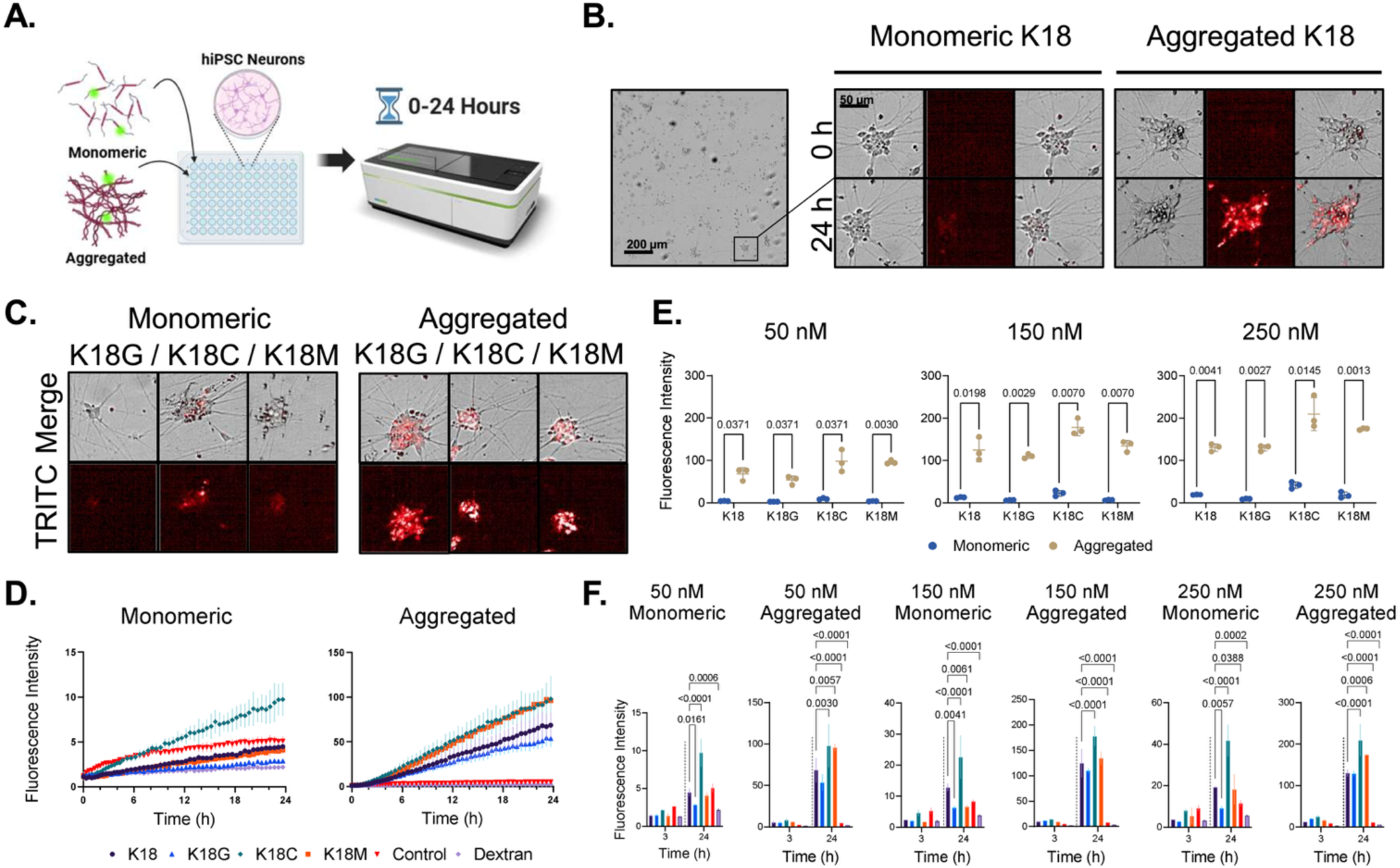
Quantification of cellular uptake of K18 proteoforms. **(A)** Schematic of the experimental workflow used to quantify cellular uptake. **(B)** Representative images of monomeric and aggregated pHrodo-labeled K18 uptake at 0 and 24 h. Left: brightfield; middle: TRITC; right: merge. **(C)** Representative images of K18 uptake at 24 h for different glycoforms. **(D)** Kinetic summary of K18 uptake at 50 nM (mean ± SD). Dextran is a pHrodo-labeled dextran control. **(E)** Comparison of monomeric versus aggregated K18 uptake at 24 h (mean ± SD; unpaired Welch’s t-test; n = 3). **(F)** Quantified uptake of Tau proteoforms (mean ± SD; two-way ANOVA with Dunnett’s multiple comparisons between glycoforms; n = 3).

Live-cell pHrodo assays with 2N4R Tau glycoproteins also revealed dose- and proteoform-dependent differences in the uptake/acidic accumulation (Figure 5). Across conditions, aggregated species achieved higher plateaus than soluble counterparts (e.g., WT at 50 nM: 14.76 vs 8.91 a.u., +65.7%), while effects on half-times were proteoform-specific. Site-selective glycosylation modulated uptake in a strikingly asymmetric manner: at 150 nM, high-mannose N410M exhibited the largest aggregate plateau (45.59 a.u., +81% vs WT-aggregated) and a faster soluble half-time (Δt_50_ = −0.50 h vs WT-Sol); N410G likewise increased aggregate plateaus (+36% vs WT-aggregated). In contrast, N359M markedly reduced soluble plateaus (−66% vs WT-Sol) and slowed soluble uptake (e.g., +1.29 h at 150 nM), with aggregated vs soluble t_50_ shortened at 150 nM. These data support a site-specific glycan effect in which modifications near the microtubule-binding repeat (Asn359) suppress uptake/accumulation, whereas a C-terminal site (Asn410) enhances it, especially for aggregates. We also performed preliminary co-localization studies with early and late endosome markers as well as a lysosome marker for K18 and 2N4R Tau proteins, and found no significant differences between the glycan composition or their position and co-localization with the organelle markers (for details, see the supporting materials).

**Figure 5.**
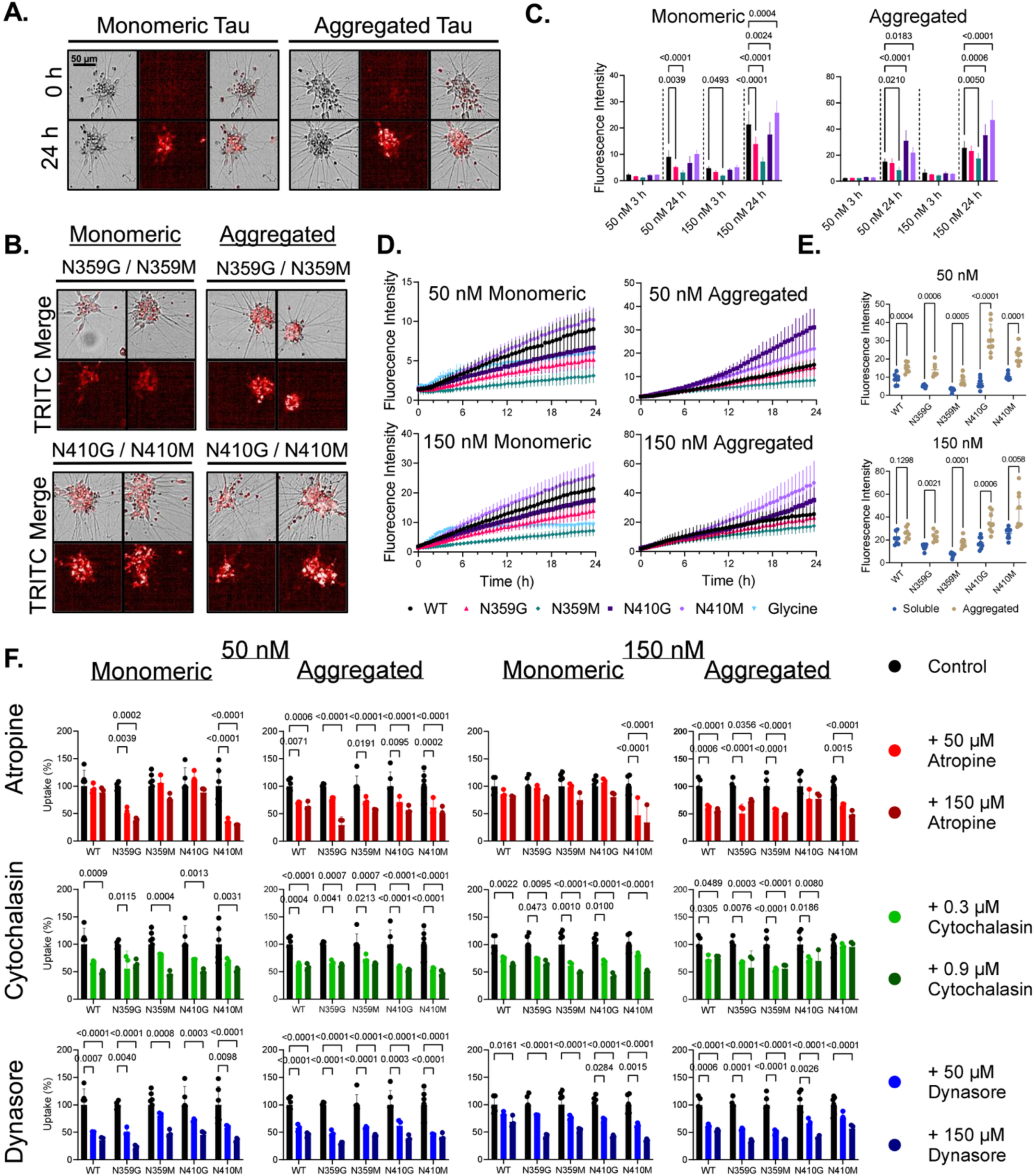
Quantification of cellular uptake and inhibitor sensitivity of Tau proteoforms. **(A)** Representative images of monomeric and aggregated pHrodo-labeled Tau uptake at 0 and 24 h. Left: brightfield; middle: TRITC; right: merge. **(B)** Representative images of Tau uptake at 24 h for different glycoforms. **(C)** Quantified uptake of Tau proteoforms (mean ± SD; two-way ANOVA with Dunnett’s multiple comparisons between proteoforms; n = 9). **(D)** Kinetic summary of Tau uptake (mean ± SD). Glycine is a pHrodo-labeled glycine as a dye control. **(E)** Comparison of monomeric versus aggregated Tau uptake at 24 h, normalized to the no-inhibitor control (100% uptake) (mean ± SD; unpaired Welch’s t-test; n = 9). **(F)** Tau uptake at 24 h in the presence of uptake inhibitors (mean ± SD; two-way ANOVA with Dunnett’s multiple comparisons; n = 9 for control, n = 3 per inhibitor).

We next profiled how three mechanistic inhibitors: atropine (muscarinic receptor blocker; 50 or 150 µM),^38, 39^ dynasore (dynamin inhibitor; 50 or 150 µM),^40–42^ and cytochalasin D (CytoD, actin/macropinocytosis inhibitor; 0.3 or 0.9 µM)^42^ influence neuronal uptake of full-length 2N4R Tau proteoforms (Figure 5F). Each condition was normalized to its matched no-inhibitor control (% of control, “100” = untreated), allowing direct quantification of % inhibition (= 100 − % of control) within proteoforms across aggregation states and protein concentrations. Three general features emerged: first, strongest effects of atropine concentrate in the monomeric state and are heavily proteoform-dependent, with the *N*-glycan at Asn410 conferring striking sensitivity. Second, dynasore robustly suppresses uptake across multiple proteoforms, especially for N359G and N410M/N359M at higher Tau doses, implicating a substantive dynamin-dependent endocytic component that is modulated by glycan site. Third, CytoD produces broad but pattern-specific inhibition consistent with actin-driven macropinocytosis yet reveals dose/aggregation asymmetries (including one notable resistance phenotype for aggregated N410M at low CytoD).

Atropine produced a clear, dose-dependent suppression of uptake that was strongest in the monomeric state and was markedly proteoform-selective. The most responsive proteoform was soluble N410M. At 50 nM Tau, N410M fell to 36.2% of control at 50 µM atropine to 29.6% at 150 µM, whereas soluble WT changed modestly from 95.8% to 88.5%. At 150 nM tTu, soluble N410M decreased from 47.0% to 34.1% as atropine increased from 50 to 150 µM; soluble N410G, which showed little or no inhibition at 50 µM (106.8%), dropped to 80.4% at 150 µM. WT in the same setting moved from 85.9% to 77.7%. Among aggregates, atropine effects were generally smaller, though N359M at 150 nM demonstrated a clearer dose step (54.6% to 47.4% as atropine increased), compared with WT (61.8% to 55.1%). Within the soluble-150 nM family, Holm-corrected tests supported significant dose-escalation effects for N410M and N410G and confirmed that these Asn410 variants were more inhibited than WT at the higher dose. Together, these results indicate a muscarinic-linked component of soluble Tau uptake that is amplified by Asn410 glycosylation and strengthened by raising atropine concentration from 50 to 150 µM.

Cytochalasin D revealed dose dependence of actin polymerization/macropinocytosis: increasing the concentration from 0.3 to 0.9 µM consistently raised % inhibition in soluble 150 nM N410G, N410M, and N359M. Of note, a low-dose anomaly and relative resistance of N410M aggregates at 150 nM to 0.3 µM cytochalasin D was maintained at 0.9 µM, where inhibition reached non-significant values. These data are consistent with a threshold-dependent reliance on actin remodeling for aggregate uptake and with increased actin involvement for select soluble proteoforms at high Tau concentrations.

Dynamin dependence also scaled strongly with dose and, like atropine, displayed specific peaks by proteoform and ligand load. For soluble 150 nM glycoprotein, N410M decreased from 62.5 ± 2.2% of control at 50 µM dynasore to 35.1% at 150 µM; N359M fell from 76.3% to 53.4%. Under the same conditions, soluble WT changed from 82.8% to 68.8%. For aggregated 150 nM Tau, N359M dropped from approximately 50% to 36.8%, outpacing the WT decrease (61.7% to 53.1%). Dose-escalation tests within dynasore families supported significant high-dose effects for soluble N410M and aggregated N359M at 150 nM, and these conditions were more inhibited than WT at 150 µM. Thus, elevating dynasore from 50 to 150 µM unmasks additional dynamin-sensitive uptake for the Asn410 high-mannose soluble proteoform at high Tau and for the Asn359 high-mannose aggregate at high Tau.

Comparisons across inhibitors and states clarify how aggregation and glycan position reweight entry pathways. Under atropine, the soluble state, especially Asn410 glycoforms, was more sensitive than the aggregated state, and sensitivity increased from 50 to 150 µM. Under dynasore, aggregated N359M at high Tau became more dynamin-dependent than its soluble counterpart, while soluble N410M at higher concentration was strongly dynamin-sensitive. Raising Tau concentration from 50 to 150 nM generally magnified dynasore and cytochalasin D effects, consistent with greater engagement of capacity-limited and actin-intensive routes when ligand load is increased.

## DISCUSSION

Glycosylation is one of the most prevalent and complex post-translational modifications in the human brain, where it plays critical roles in protein folding, trafficking, stability, and intercellular communication.^43, 44^ In the context of AD, increasing evidence suggests that aberrant glycosylation, particularly in the form of altered *N*-linked glycan profiles, may contribute to disease pathogenesis.^18, 45, 46^ Comparative analyses of the *N*-glycome in healthy and AD brains have revealed significant differences in glycan structure and composition that may influence the behavior of key proteins involved in neurodegeneration.^8, 47^ For proteins such as neurotransmitter receptors,^48, 49^ ion channels,^50^ adhesion molecules,^51, 52^ and immune receptors,^53^ proper *N*-glycosylation is required for normal function. Tau, however, presents an intriguing case. Several studies have identified *N*-glycosylated forms of Tau in AD brains, typically in association with hyperphosphorylation and aggregation.^8–12^ This aberrant glycosylation appears to be linked to disease-specific conformations and may contribute to misfolding and resistance to degradation.

Despite the absence of a classical secretion signal, Tau is actively released from neurons through a variety of non-canonical mechanisms. Once in the extracellular space, Tau can be taken up by neighboring cells through several routes. One major mechanism involves endocytosis, including clathrin-mediated endocytosis and macropinocytosis. Cell-surface HSPGs facilitate the binding and uptake of Tau aggregates, and the specific sulfation patterns of these glycans influence the efficiency of internalization. Our quantitative BLI measurements reveal two mechanistic binding regimes: heparin/HSPG-like interactions with full-length 2N4R Tau are avidity-dominated - strong electrostatic attraction to clustered basic patches yields low-nanomolar K_D_ and slow off-rates (multi-minute residence), consistent with prior SPR/BLI reports that emphasize sulfate patterning and multivalency as key determinants of affinity and trafficking. Site-specific *N*-glycans at Asn359 or Asn410 reduce this avidity in 2N4R Tau, most likely by sterically shielding charge, altering local hydration, and modestly shifting the conformational ensemble, thereby dampening HSPG engagement at the cell surface. The smaller glycan exerts a more pronounced reduction in affinity, suggesting that compact *N*-linked structures can more effectively shield local charge and sterically interfere with heparin engagement. By contrast, the bulkier high-mannose glycans appear to maintain Tau in a more conformationally flexible state that can better accommodate heparin interactions, attenuating the penalty in K_D_. In this view, larger glycans preserve a degree of binding-compatible configurational freedom, whereas smaller glycans strike a different balance between steric size and local dynamics that renders Tau less susceptible to high-avidity heparin engagement. Second, RNA interactions (A40/C40/U40) fall in a nanomolar–sub-micromolar K_D_ range with faster kinetics. Here, small GlcNAc at Asn359/Asn410 can enhance binding (notably to U40), plausibly by loosening long-range intramolecular contacts (e.g., PRR–MTBR “paperclip”) and exposing nucleic-acid-interacting patches without replicating the dense electrostatic avidity required for maximal heparin binding.

In parallel with the uptake and binding data, our cross-seeding kinetics show that site-specific *N*-glycosylation reweights templating compatibility between Tau proteoforms in a position- and glycan-dependent manner. For K18 and K18M monomers seeded with K18-family fibrils, small N359 glycans on the seed preserved or even enhanced nucleation, whereas high-mannose at N359 weakened early nucleation despite steep late-phase growth and large endpoints. With full-length 2N4R seeds, N410 glyco-variants are the most potent accelerants of K18 (N410G ≥ N410M > N359G ≳ Tau > N359M). Mechanistically, these seeding hierarchies align with our heparin/HSPG and receptor-mediated uptake regimes (*vide infra*): Asn359 (within/adjacent to the MTBR) likely sterically/hydrationally dampens nucleation-competent docking and avidity, slowing t₅₀ even when elongation remains fast, whereas Asn410 (outside the MTBR) preserves β-sheet templating faces and boosts cross-seeding.

Additionally, receptor-mediated uptake contributes to Tau transmission. The robust dynasore sensitivity for both soluble and aggregate conditions, dose-responsive CytoD suppression, and the unusually large atropine sensitivity of soluble Asn410 glyco variants, fit into a framework in which extracellular Tau uses receptor-mediated, dynamin-dependent endocytosis together with actin-dependent macropinocytic/bulk processes, with the balance shifted by aggregation state and by the site of *N*-glycosylation on tau. In human iPSC-derived neurons both monomeric and aggregated Tau enter efficiently. Aggregated Tau entry is largely dynamin-dependent and actin-independent, and monomeric Tau uses two routes: a rapid, dynamin-dependent phase and a slower, actin-dependent phase.^54^ LRP1 as a primary receptor for monomeric/oligomeric Tau and LRRK2 acts as a key regulator of monomeric (but not fibrillar) Tau uptake validating dynamin/PI3K/v-ATPase and broader endolysosomal machinery as central nodes. Our dynasore results are concordant and add a dose dimension. Raising dynasore from 50 to 150 µM augments inhibition in soluble N410M Tau and in aggregated N359M tau, and these high-dose conditions are more inhibited than WT. That mirrors recent conclusion that aggregated Tau in human neurons is “almost entirely” dynamin-dependent,^54^ and genetic evidence of a broad dynamin/PI3K/v-ATPase scaffold downstream of the initial receptor engagement. Our observation that N359M aggregates show the steepest dynasore dose-response at high Tau load fits a receptor-usage shift away from LRP1 toward non-LRP1 endocytic routes that still rely on dynamin and the endolysosomal axis. The CytoD titration adds nuance to this mechanistic picture. Little effect of actin disruption on aggregate entry was reported in human neurons over 3-4 h, whereas we see clear dose-dependence at 24 h (0.9 µM > 0.3 µM) for several aggregates (notably N359M and N410M) and for soluble Asn410/Asn359 variants at high Tau. Two factors plausibly reconcile the difference. First, pHrodo endpoints over 24 h aggregate entry/acidic routing emphasize cumulative trafficking more than the early 3-4 h imaging windows used before.^54^ Second, fibrillar uptake depends on glycosylation pathways and HSPG biology, and HSPGs are central for fibrils in neurons - a setting where actin-dependent macropinocytosis-like processes can contribute, particularly at higher Tau load or with glycan additions that enhance avidity/multivalency. In short, our 24 h, high-dose and glycoengineered conditions can “tip” aggregates into a measurably actin-dependent regime, without contradicting the earlier 3-4 h observation that initial aggregate internalization is dominated by dynamin.

The atropine study merges with the prior receptor-centric models. Soluble Asn410 glyco variants (N410M ≫ N410G) show marked atropine sensitivity that increases with dose and with Tau concentration, whereas aggregated phenotypes are comparatively less atropine sensitive. LRP1 is at the heart of monomeric/oligomeric uptake, while showing fibrils do not depend on LRP1 at threshold.^16^ Our atropine pattern is therefore most parsimonious with a receptor-proximal signaling axis that modulates the efficiency of LRP1-linked, dynamin-dependent uptake for soluble Tau where we see the largest atropine effects, and that is amplified by Asn410 glycosylation outside the MTBR, which preserves MTBR-mediated interactions (including LRP1 engagement) while potentiating signaling-dependent entry/routing.

Integrating these dose-aware inhibitor effects with our prior time-course kinetics supports a route-partitioning model for Tau entry into human neurons that is co-governed by aggregation state, glycan position, inhibitor dose, and Tau load. Glycan position offers a simple molecular rationale for our proteoform specificity. Monomeric entry is LRP1-mediated while fibrillar entry uses other receptors and HSPG-dependent mechanisms, with strong endolysosomal/PI3K/v-ATPase dependencies downstream. An Asn359 glycan, within/near the MTBR, is ideally placed to alter the LRP1-binding face and thereby dampen soluble LRP1 engagement, consistent with our smaller soluble plateaus for N359M in no-inhibitor conditions and its greater reliance on dynamin/actin once uptake is challenged, especially for aggregates at high dose. By contrast, Asn410 lies outside the MTBR and would be expected to preserve the MTBR surface while changing multivalency/spacing, providing a mechanistic route to the striking atropine and dynasore dose-responses we observe for soluble N410M. Involvement of other receptors that interact with high-mannose glycans and facilitate the uptake cannot be excluded at the point,^55^ and likely can contribute more significantly to the Asn410 modifications rather than to the glycosylation at Asn359. Additionally, the recent LRRK2 findings help contextualize the dose changes without perturbing LRRK2 directly.^56^ LRRK2 kinase inhibition depresses uptake of both monomeric and fibrillar Tau in human neurons, while genetic modulation of LRRK2 remodels endolysosomal traffic and surface levels of LRP1. Given that dynamin, PI3K class III and v-ATPase all emerged in the prior study, it is unsurprising that high-dose dynasore and CytoD in our assays unmask additional inhibition in proteoforms/states that ride most heavily on these pathways at 24 h. The modest divergences from earlier reports, most notably the actin-independence of aggregates in early time windows, are accounted for by assay horizon (24 h vs 3-4 h), proteoform chemistry (site-specific *N*-glycans), and ligand dose.^42, 56^ Together with prior receptor and genetic data, our findings support the notion that site-specific *N*-glycosylation can be deployed as a molecular “dial” to re-weight Tau uptake pathways, and that dose-tunable pharmacology targeting muscarinic receptors, dynamin, and actin remodeling can steer Tau trafficking toward more degradative, non-seeding fates.

In sum, our inhibitor and binding data support a model in which soluble Asn410 glyco-variants preferentially exploit a muscarinic-tunable, LRP1/dynamin-dominated route whose efficiency scales with inhibitor dose and ligand load, whereas Asn359 glycosylation biases aggregates toward a dynamin/actin-intensive itinerary whose macropinocytic component becomes increasingly visible at higher CytoD and higher Tau.

## METHODS

### Glycoprotein Preparation

*N*-Glycosylated Tau proteins were produced and characterized as described previously.^25^ The full-length construct (“2N4R Tau”) corresponds to microtubule-associated protein Tau isoform 2 (MAPT; NCBI RefSeq NP_005901.2). The K18 construct comprises residues 244–372 of the 2N4R isoform. Glycoforms were specified by site and glycan composition, as shown in Figure 1. For 2N4R Tau, a single *N*-acetylglucosamine (GlcNAc) was installed at Asn359 (N359G) or Asn410 (N410G), and a high-mannose glycan was installed at Asn359 (N359M) or Asn410 (N410M). For K18 Tau, glycosylation was restricted to Asn359 and included GlcNAc (K18G), chitobiose (GlcNAc–GlcNAc; K18C), or a high-mannose glycan (K18M).

### Seeded Aggregation of K18 Monomers

Tau fibrils were prepared as described before.^25^ A 0.65 µL Eppendorf tube was charged with the 50 µL aggregation assay mixture (or 25 µL for 2N4R Tau fibrils) at the final concentration listed. The buffers and salts were added first, followed by protein, heparin, and the seeds were added last. A black, 384-well, non-binding microplate (Greiner BioOne, PS, F Bottom, small volume, HiBase, #784900) was filled with the aggregation assay mixture (3 wells with 15 µL per well for K18 seeds, or 2 wells with 10 µL per well for 2N4R Tau seeds), the plate was sealed with polyester adhesive film (VWR #89134-430) and incubated at 37 ⁰C in the microplate reader. The data from the three wells were averaged, and the experiment was repeated a total of three times. The data was averaged then normalized, and the error bars on the graphs are reported as the standard error of the mean.

Stock Solutions: 25 mM HEPES (pH 7.4); 100 mM DTT in 25 mM HEPES (pH 7.4); 1 M NaCl in 25 mM HEPES (pH 7.4); 110 mM heparin sodium salt from porcine intestinal mucosa (Sigma H4784, 18 kDa average molar weight, 1 mg/mL = 55 µM) in 25 mM HEPES (pH 7.4); 165 µM Thioflavin T in 25 mM HEPES (pH 7.4); 200 µM K18 in 25 mM HEPES (pH 7.4); 200 µM K18M in 25 mM HEPES (pH 7.4); 50 µM K18-Heparin fibrils in assay buffer; 20 µM Tau-Heparin fibrils in assay buffer; 43 µM Tau PFFs in PBS, Acro Biosystems H5115.

Final Conditions: 25 µM K18, 5 µM heparin, 2.5 µM K18-heparin fibrils, 10 µM thioflavin T, 10 mM DTT, 100 mM NaCl, 25 mM HEPES (pH 7.4), 37 ⁰C; 25 µM K18M, 5 µM heparin, 2.5 µM K18-heparin fibrils, 10 µM thioflavin T, 10 mM DTT, 100 mM NaCl, 25 mM HEPES (pH 7.4), 37 ⁰C; 25 µM K18, 5 µM heparin, 2.5 µM Tau-heparin fibrils, 10 µM thioflavin T, 10 mM DTT, 100 mM NaCl, 25 mM HEPES (pH 7.4), 37 ⁰C.

Microplate Settings for K18: wavelength: excitation at 440 nm, emission at 480 nm. PMT gain: 500 V, read 9 mm from the top of the plate. Integration time: 400 ms. Timing: Read every 10 minutes over 72 h; shake for 30 s before the first read, and then no shaking.

Kinetic accelerator factors parameters were extracted with the Prism software (GraphPad) by one phase association nonlinear regression of normalized ThT signal. Acceleration factor with 95% CI was calculated by AF_t50_ = t_50_(unseeded)/t_50_(seed) with t_50_ = half-time (h).

### Biolayer Interferometry (BLI)

The biolayer interferometry experiments were performed on a Sartorius Octet N1 system. Octet High Precision Streptavidin (SAX) biosensors were hydrated with 1× PBS (pH 7.4) for 10 minutes at 23 °C. Ligand was immobilized by incubating 4 µM heparin-biotin sodium salt conjugate (Sigma #B9806), or 5’-biotinylated A(40), C(40), or U(40) (Genescript) for 1 h. The sensors were washed with PBS for 2 minutes prior to being dipped into a concentration range of Tau or K18. Binding kinetics were monitored and analyzed using the Octet system. Experiments were performed in triplicate and recorded as the SD. The stock solutions used: heparin-biotin sodium salt, 5’-biotinylated A(40) RNA, 5’-biotinylated C(40) RNA, and 5’-biotinylated U(40) RNA all at 4 µM in 1× PBS (pH 7.4). Tau dilutions: 20 µM, 10 µM, 5 µM, 2.5 µM, 1.75 µM in 1× PBS pH 7.4.

### iPSC Maintenance and i^3^N Differentiation

WTC-11 iPSCs were obtained as a gift from Dr. Chris Link (CU-Boulder) and were maintained and differentiated into i^3^Neurons as described previously.^57, 58^ WTC-11 cells were cultured on Matrigel hESC-qualified matrix (Corning, #354277) and were fed daily with B8 medium (Burridge) prepared in-house. Cells were maintained at 37 °C, 5% CO₂ in a water-jacketed incubator. Passaging was performed approximately every 48 h, and colonies retained normal morphology with minimal spontaneous differentiation. For cortical neuron induction, WTC-11 cells were expanded and were seeded as single cells onto Matrigel-coated 150-mm plates (2.4 × 10^7^ cells per plate) in Induction Medium [DMEM/F12 (Gibco, #11330-032), 1× N2 Supplement (Gibco, #17502048), 1× non-essential amino acids (Gibco, #11140050), 1× GlutaMAX (Gibco, #35050-061), and 2 mg/mL doxycycline] supplemented with 10 µM Y-27632 ROCK inhibitor for ∼24 h following passaging, after which medium was changed to Induction Medium without ROCK inhibitor. On day 3, induced cells were dissociated with Accutase (Gibco, #A1110501), counted, resuspended in cryopreservation medium (90% KOSR, 10% DMSO) at 2.0 × 10⁶ cells/mL, and frozen in 1.5-mL cryovials. A large batch of pre-differentiated i3Neurons (140 aliquots) was prepared on the same date and passage number to support reproducible downstream experiments. Pre-differentiated i^3^Neurons were plated onto 96-well Pheno-µ-plates (PerkinElmer, #6055300) pre-coated with 0.05 mg/mL poly-D-lysine (Gibco, #A38904-01) followed by 0.05 mg/mL laminin (Gibco, #23017015); laminin was aspirated immediately prior to seeding. Cells were plated at 24,000 cells/well (or 12,000 cells/well for imaging) in 200 µL Cortical Neuron Maturation Medium [Neurobasal A, 1× B27 Supplement (Gibco, #17504044), 10 ng/mL BDNF (PeproTech, #450-02), 10 ng/mL NT-3 (PeproTech, #450-03), and 1 mg/mL laminin]. Half of the medium was replaced bi-weekly until use.

### Tau-Uptake Assay and Live Imaging

i^3^Neurons matured for 12 days were used in the assay. Tau proteoforms (labeled with 10% pHrodo red STP ester, amine reactive; ThermoFisher #P35369) were thawed on ice prior to use. Aggregated proteoforms were sonicated in a water-bath sonicator twice for 30 seconds (1 min rest in between). Half of the well volume (100 μL) was removed and replaced with 100 μL of freshly conditioned medium + 2× Tau-proteoforms (at the indicated concentrations) immediately before imaging. Neurons were imaged for fluorescent pHrodo-labeled (excitation at 561 nm and emission at 570-630 nm) Tau proteoforms using 10× Air, NA 0.3 in confocal optical mode on Opera Phenix System (Perkin Elmer) in a temperature- and CO_2_-controlled environment. Bright-field and fluorescence emission images were collected at 20 min intervals for the first 5 h, and at 30 min intervals until the end of 24 h interval.

Parameters for florescence intensity were set and quantified using Harmony 4.9 software (PerkinElmer). Conditions were repeated in triplicate with side-by-side wells and >4 of 9 frames per well (average of 8). Frames containing densely packed cells or irregular contaminants were excluded from processing. Fluorescence intensity within cells was determined by subtracting mean fluorescence of background from the mean intensity within areas identified as neurons. Neurons were identified by texture recognition on brightfield channel (trained on 10 images with 100 points used and insignificant changes in recognition occurred with additional training). These texture regions were then filtered by removing areas <500 px^2^.

Atropine (TCI #A0550) was dissolved in ethanol at a concentration of 69.5 mg/mL (100 mM) and stored at –20 °C. Cytochalasin D (Abcam #ab143484) was dissolved in DMSO at a concentration of 25 mg/mL (50 mM) and stored at –20 °C. Dynasore (Abcam #ab120192) was dissolved in DMSO at a concentration of 32.23 mg/mL (100 mM) and stored at –20 °C. For the Tau-uptake in the presence of inhibitors, half of the well volume (100 µL) was removed and replaced with 50 µL of freshly conditioned medium and 3× inhibitor concentration (or vehicle control) 1 h prior to adding 50 µL of freshly conditioned medium plus 1× inhibitor concentration (or vehicle control) and 4× Tau-proteoforms (at the indicated concentrations) immediately before imaging.

Data was analyzed with the Prism software (GraphPad). Comparison between monomeric and aggregated Tau was performed by unpaired Welch’s t test (n=3 K18, n=9 Tau) with Holm-Sidak correction for multiple comparisons. Comparison between proteoforms was performed by 2-way ANOVA with Dunnet multiple comparisons (n=3 K18, n=9 Tau). Comparisons between inhibitor concentrations were performed by 2-way ANOVA with Dunnet multiple comparisons (n=9 uninhibited, n=3 with inhibitors).

### Bioreporter Cell Seeding

HEK293T bioreporter cells stably expressing CFP/YFP K18 were obtained from ATCC (#CRL-3275) and maintained in Dulbecco’s modified Eagle’s medium (DMEM, high glucose) supplemented with GlutaMAX™ and 10% fetal bovine serum (FBS) at 37 °C in a humidified incubator with 5% CO₂. For seeding experiments, cells were plated 24 h prior to transfection at 50,000 cells per well in 24-well plates in growth medium. At ∼60% confluency, cells were transfected with a mixture of proteoform stocks and Lipofectamine 3000 prepared in Opti-MEM (total transfection volume, 50 µL per well). The transfection mixture was incubated at room temperature for 20 min before being added to the cells, resulting in a final protein concentration of 100 nM after dilution into the well volume. Cells were incubated with the transfection mixture for 24 h at 37 °C, detached with TrypLE, pelleted by centrifugation, and resuspended in ice-cold phosphate-buffered saline (PBS). Single-cell suspensions were passed through 0.22 µm filters into flow cytometry tubes (BD FACSCelesta). FRET signals were acquired on a BD FACSCelesta flow cytometer using instrument settings described previously.^27^ For each technical replicate, 10,000 events were collected. Nine technical replicates per condition and three biological replicates were analyzed. Data were processed using FlowJo v10.

### Statistical Analysis

Data was analyzed and plotted with the Prism software (GraphPad) as described above.

## ACKNOWLEDGEMENTS

This work was supported by the National Institute on Aging (RF1AG079294 and R01AG087295) and NIH/CU Molecular Biophysics Program and NIH Biophysics Training Grant (T32GM145437). The imaging work was performed at the BioFrontiers Institute’s Advanced Light Microscopy Core (RRID: SCR_018302). The Revvity Opera Phenix was supported by NIH (S10OD025072). This data analysis and visualization work were performed at the BioFrontiers Institute’s Advanced Light Microscopy Core (RRID: SCR_018302). The software package Imaris was supported by NIH (S10RR026680). Parts of Figure 4 were created with biorender.com.

## Notes

### Competing Interest Statement

The authors have declared no competing interest.

