## Supplementary Materials for "*N*-Linked Glycosylation as a Driver of Tau Pathology and Neuronal Transmission"

### 1. Seeding of K18 Monomer with K18 Seeds

**Normalized Seeding of K18 Monomer with K18 Seeds**

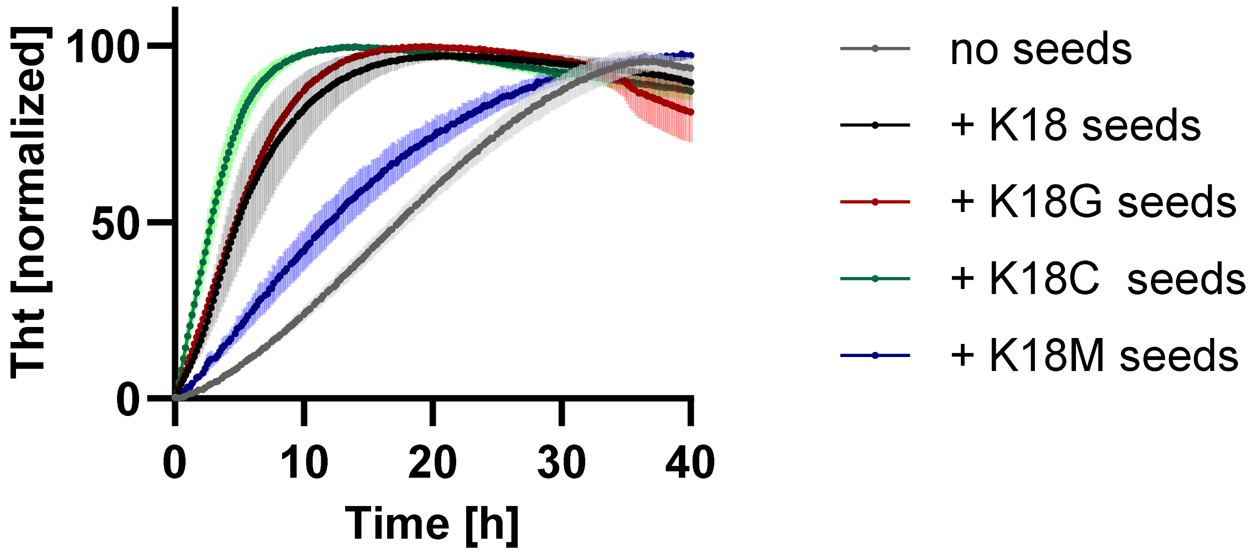

**Seeding of K18 Monomer with K18 Seeds – Individual Plots**

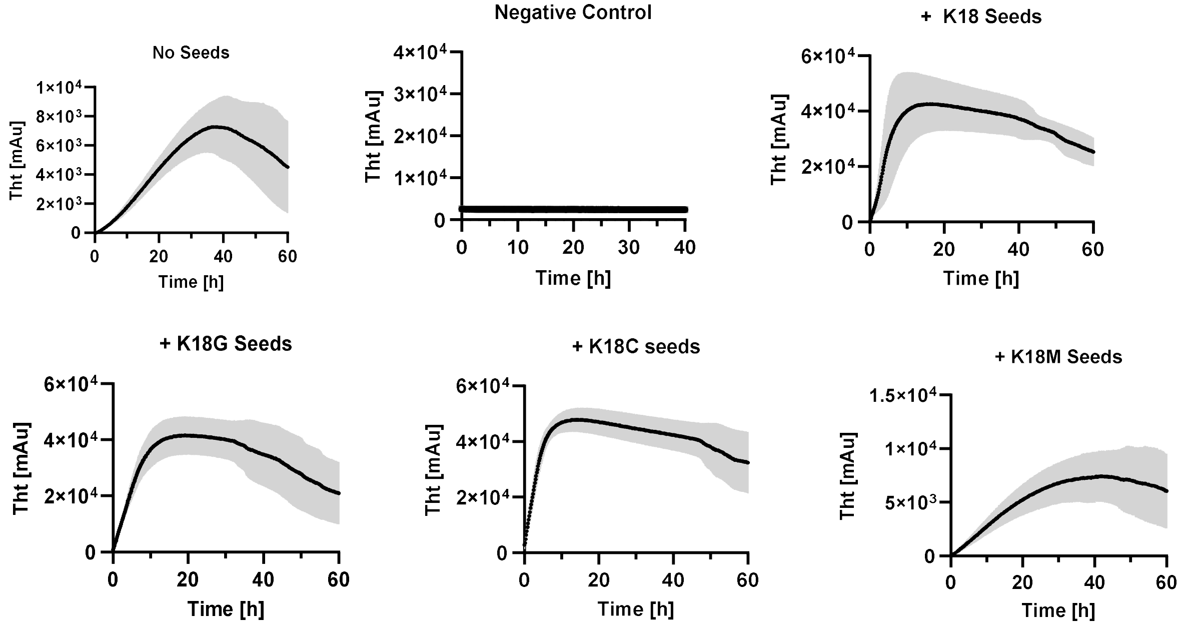

### 2. Seeding of K18M Monomer with K18 Seeds

**Normalized Seeding of K18M Monomer with K18 Seeds**

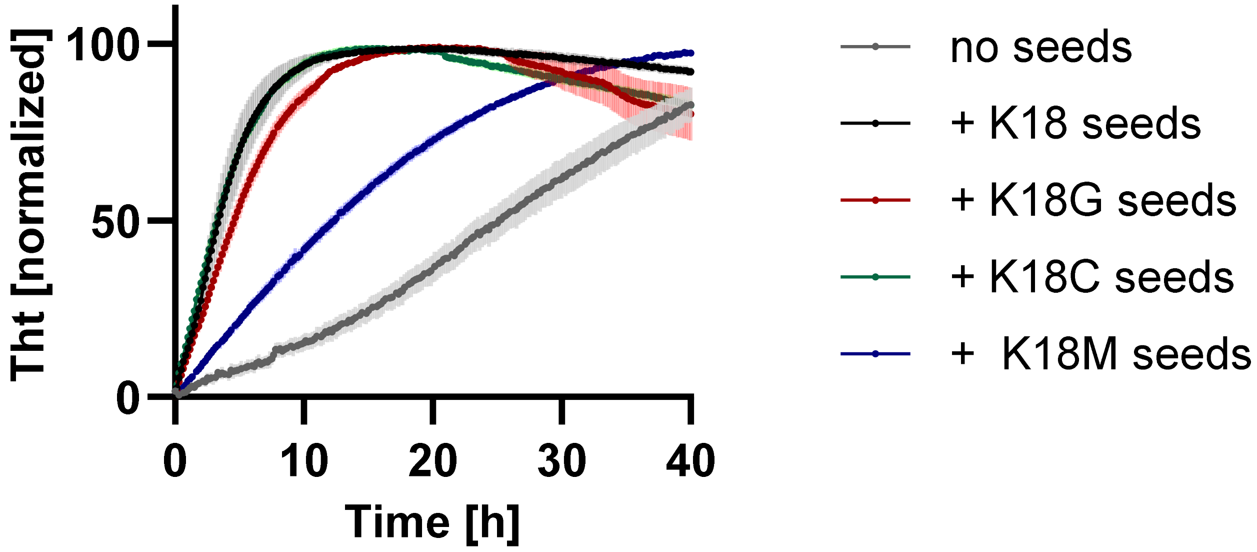

**Seeding of K18M Monomer with K18 Seeds – Individual Plots**

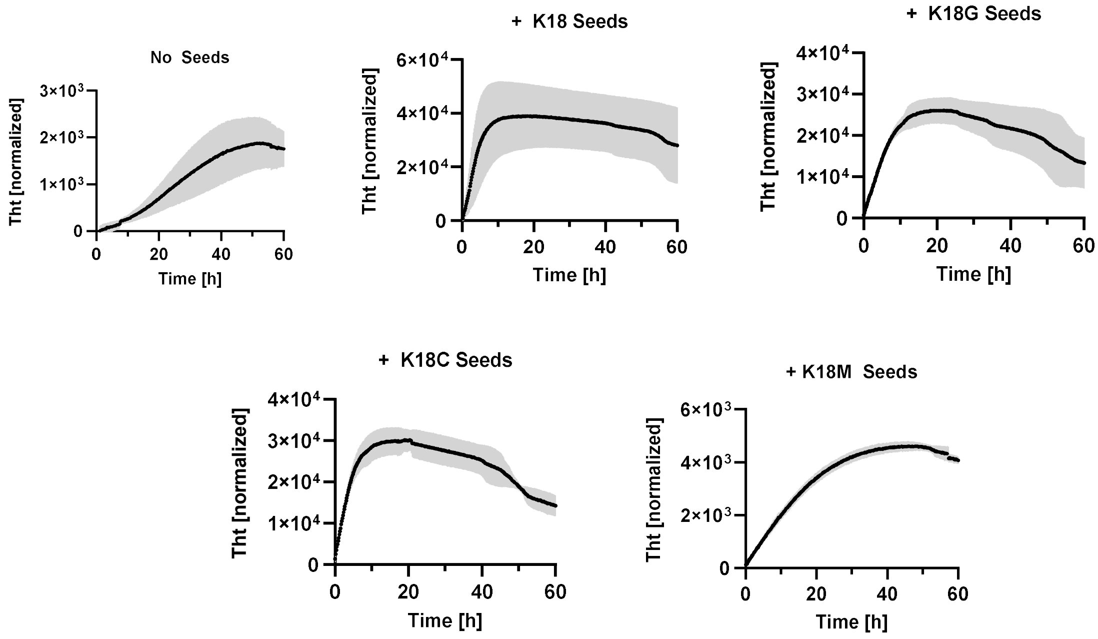

### 3. Seeding of K18 Monomer with 2N4R Tau Seeds

**Normalized Seeding of K18 Monomer with Tau Seeds**

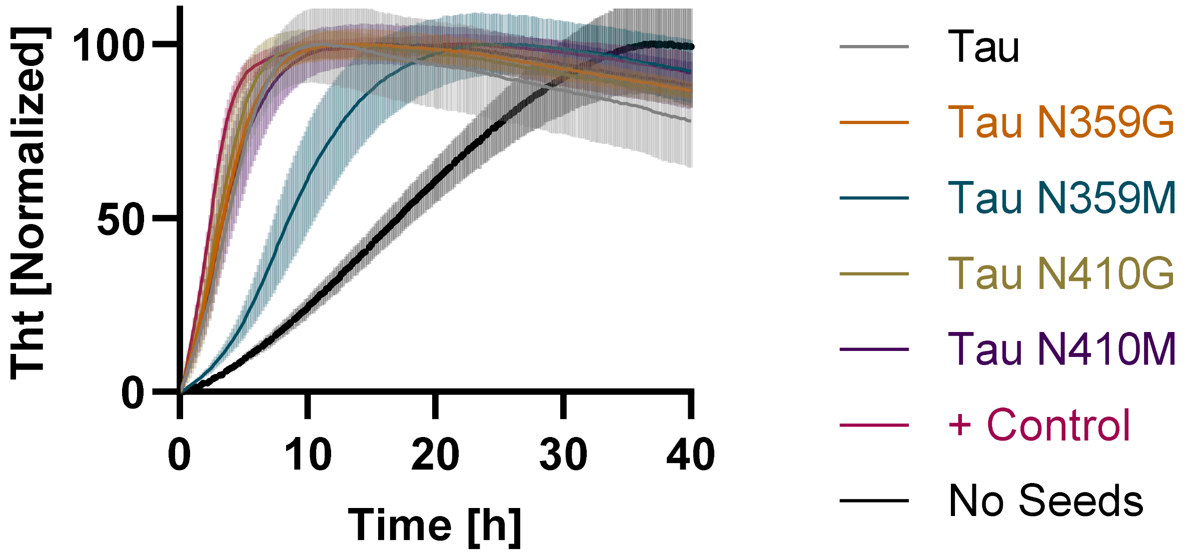

**Seeding of K18 Monomer with Tau Seeds – Individual Plots**

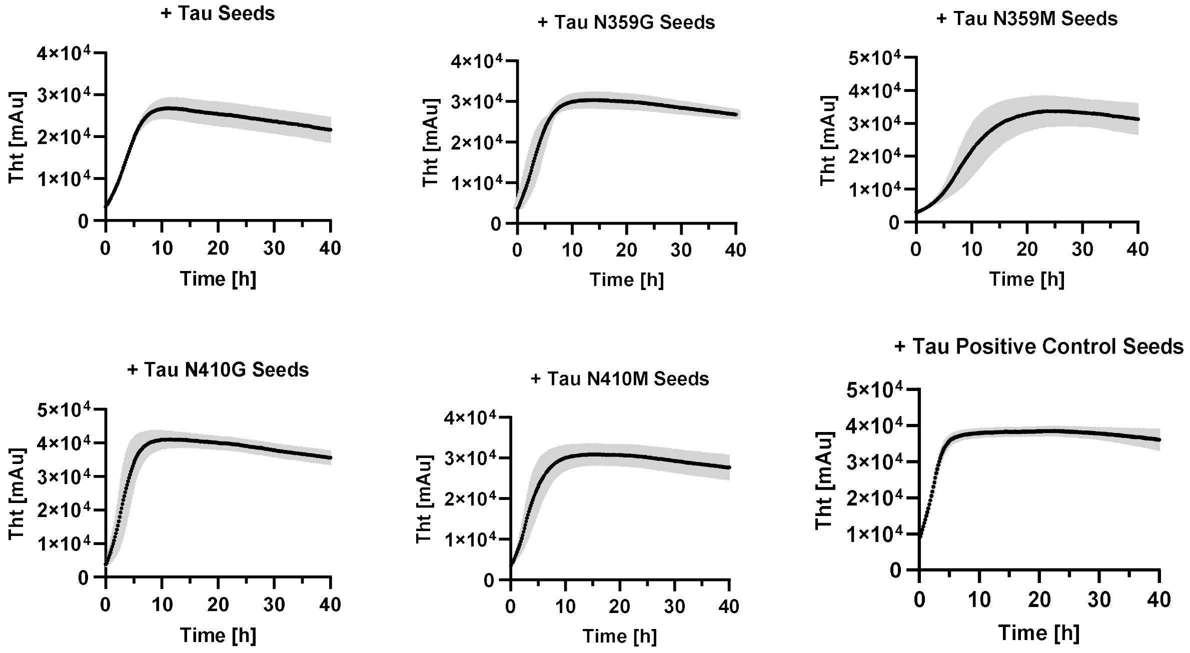

### 4. Monomeric K18 Uptake Images (50 nM)

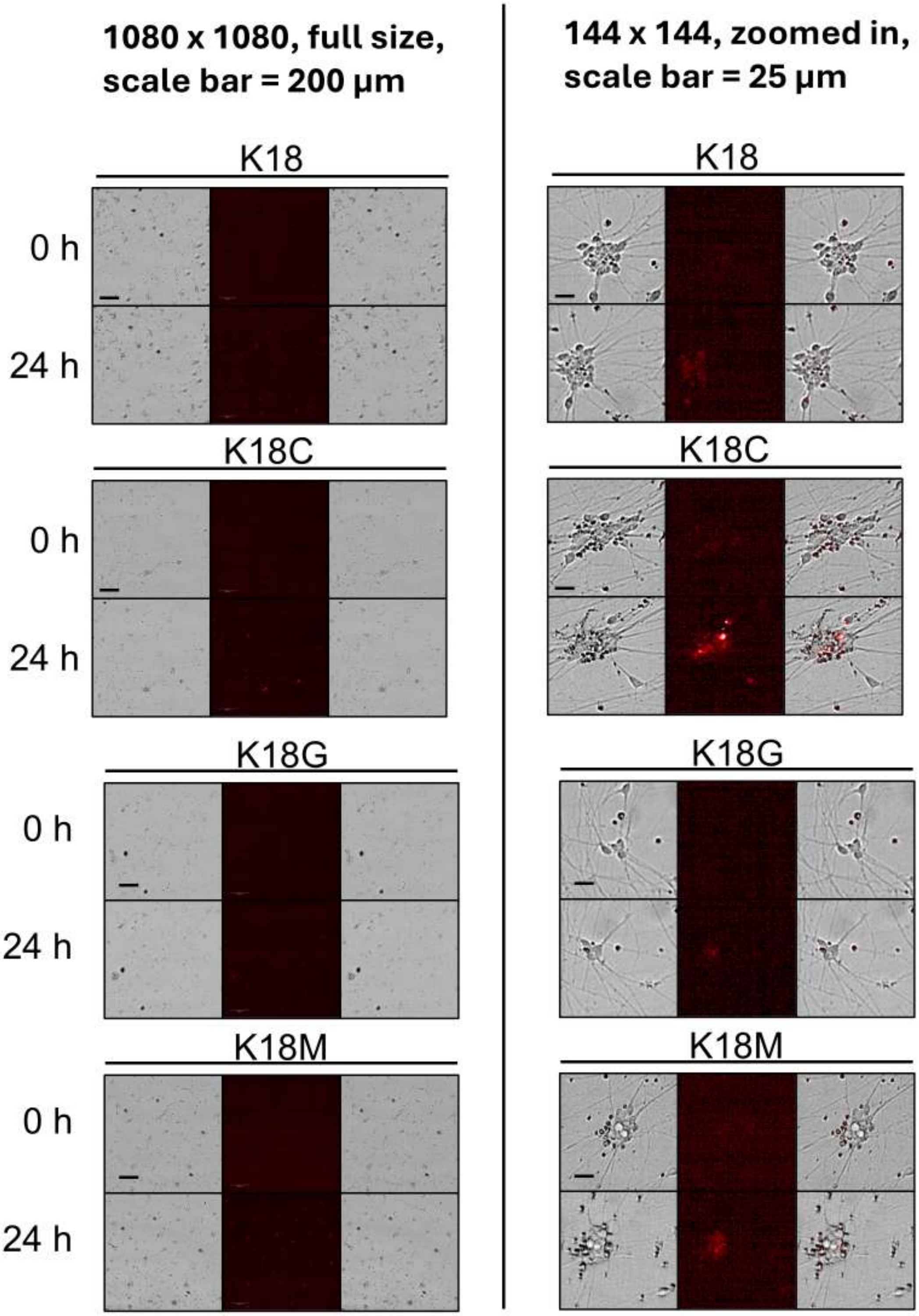

### 5. Aggregated K18 Uptake Images (50 nM)

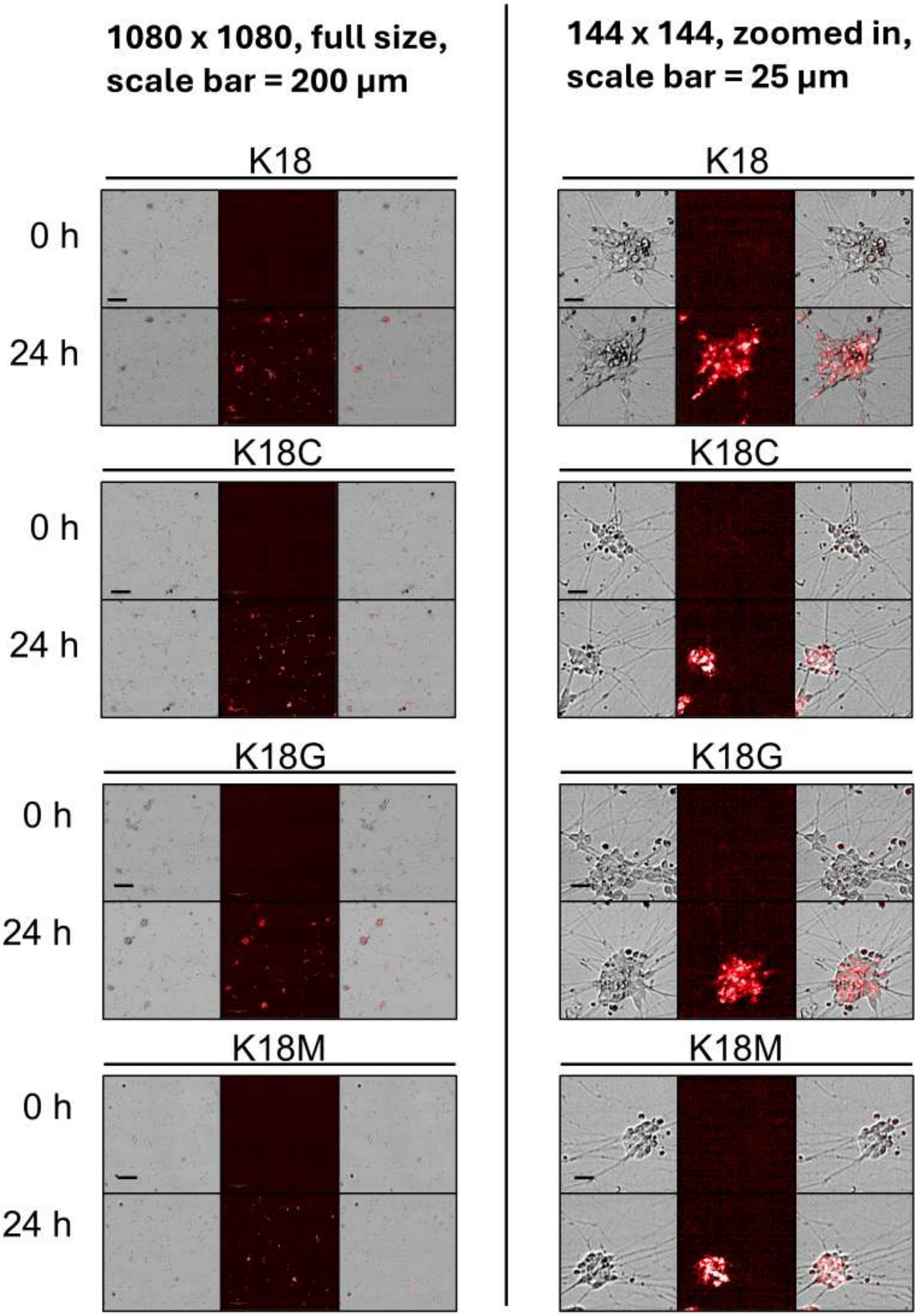

### 6. Monomeric 2N4R Tau Uptake Images (50 nM)

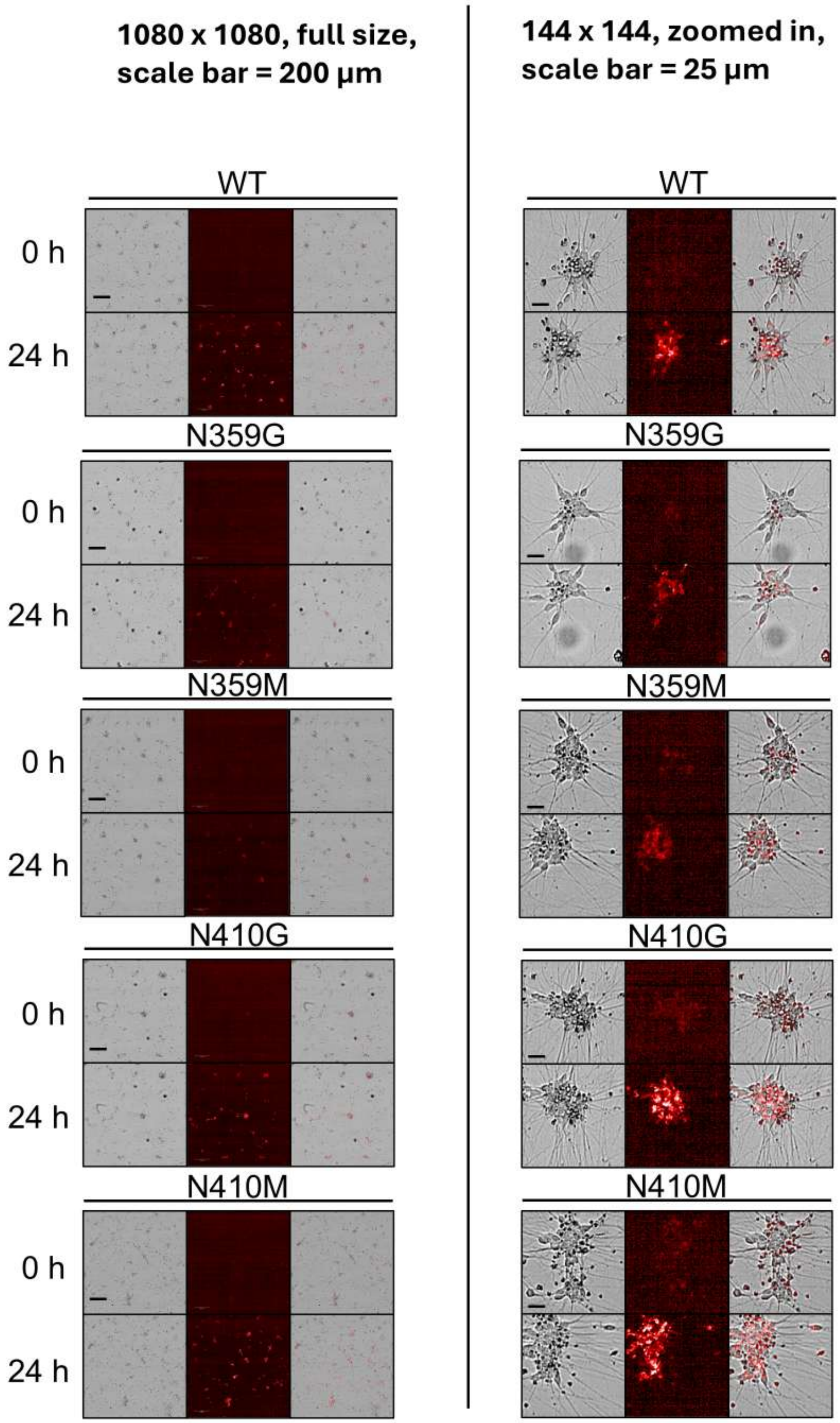

### 7. Aggregated 2N4R Tau Uptake Images (50 nM)

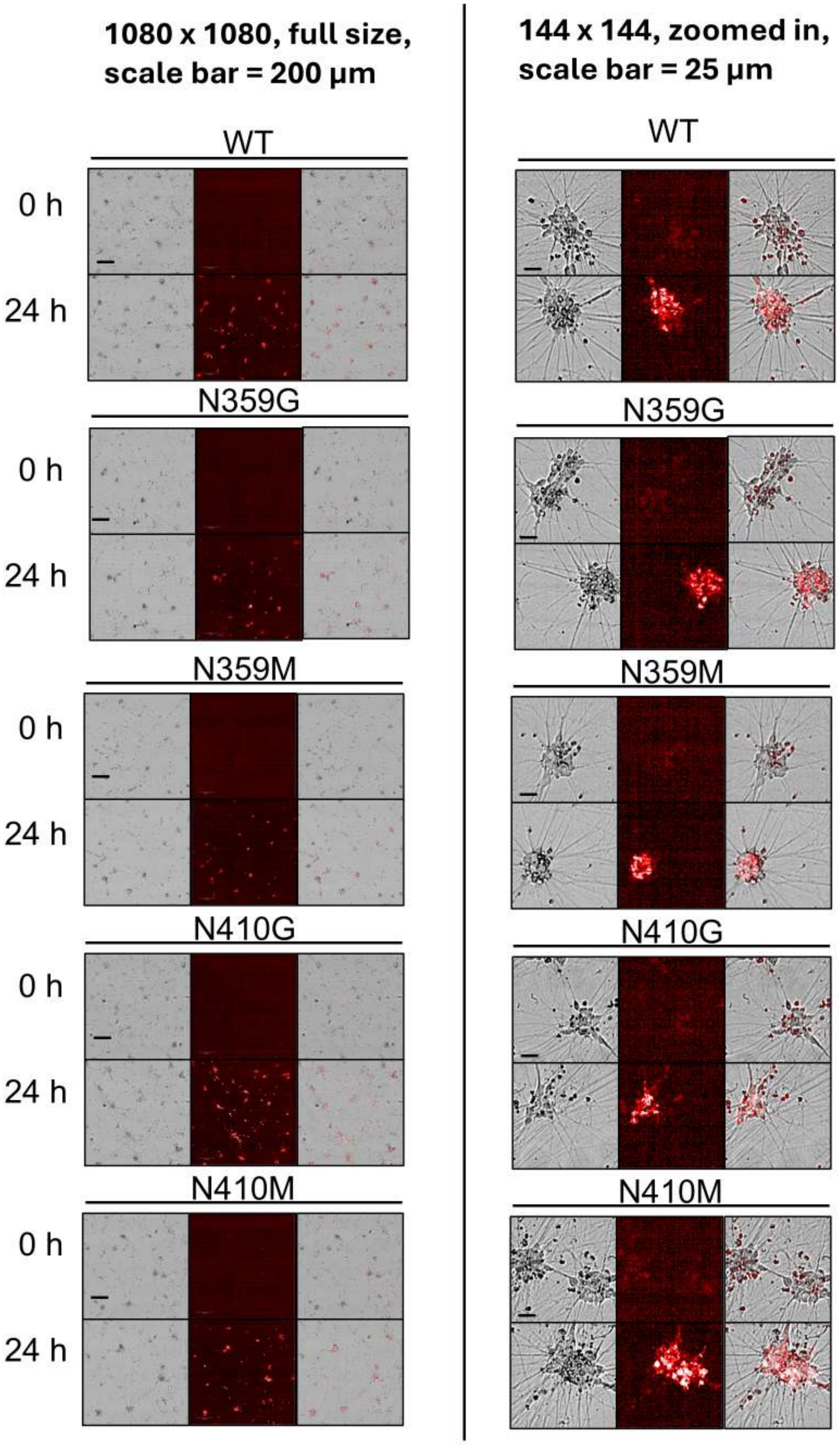

### 8. Colocalization of Monomeric 2N4R Tau with Early Endosome Tracker

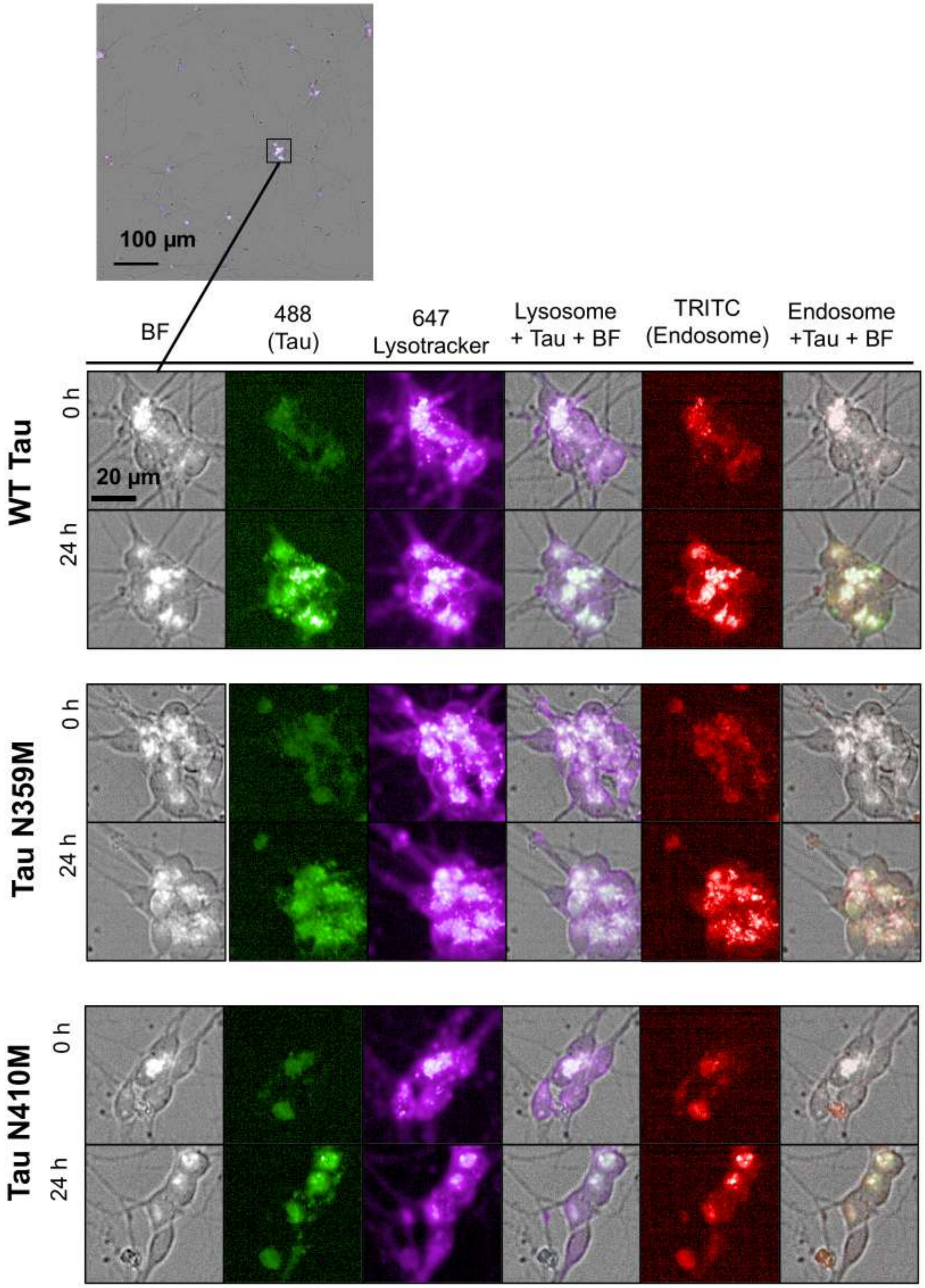

### 9. Colocalization of Aggregated 2N4R Tau with Early Endosome Tracker

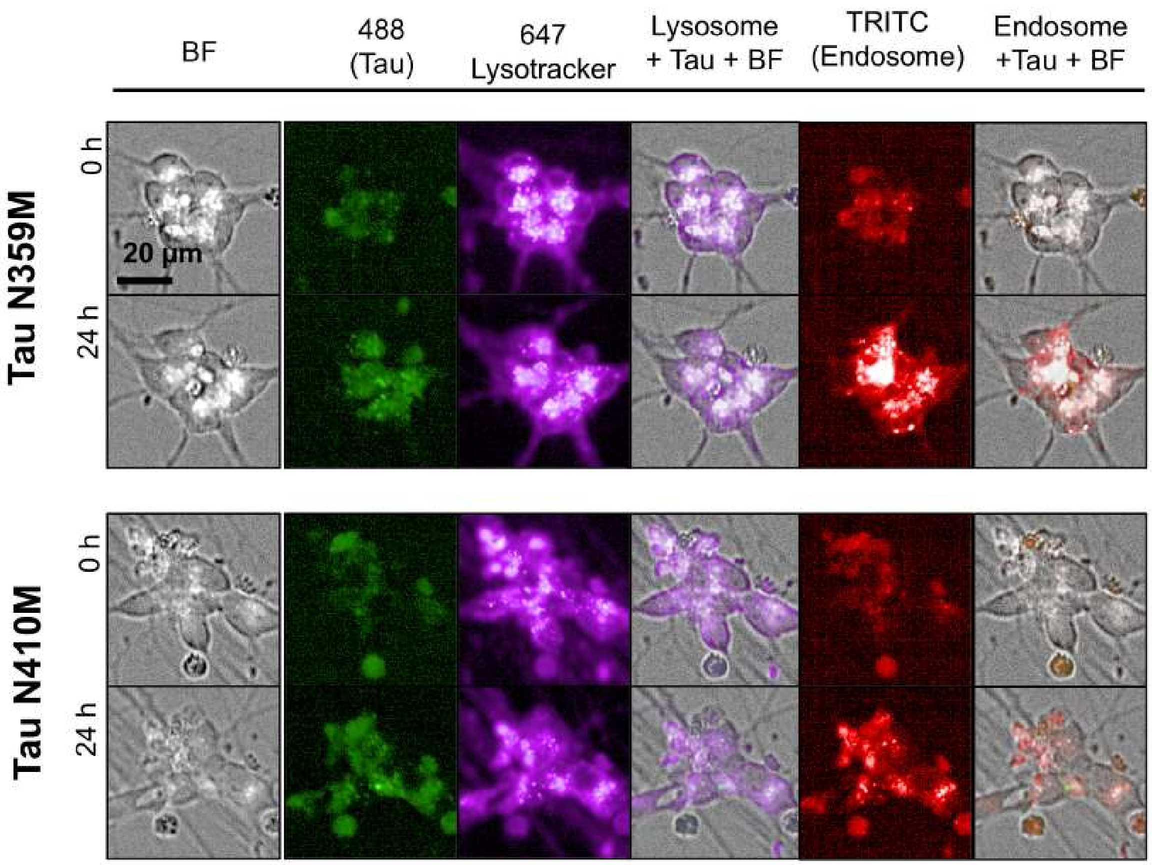

### 10. Colocalization of Monomeric K18 Tau with Early Endosome Tracker

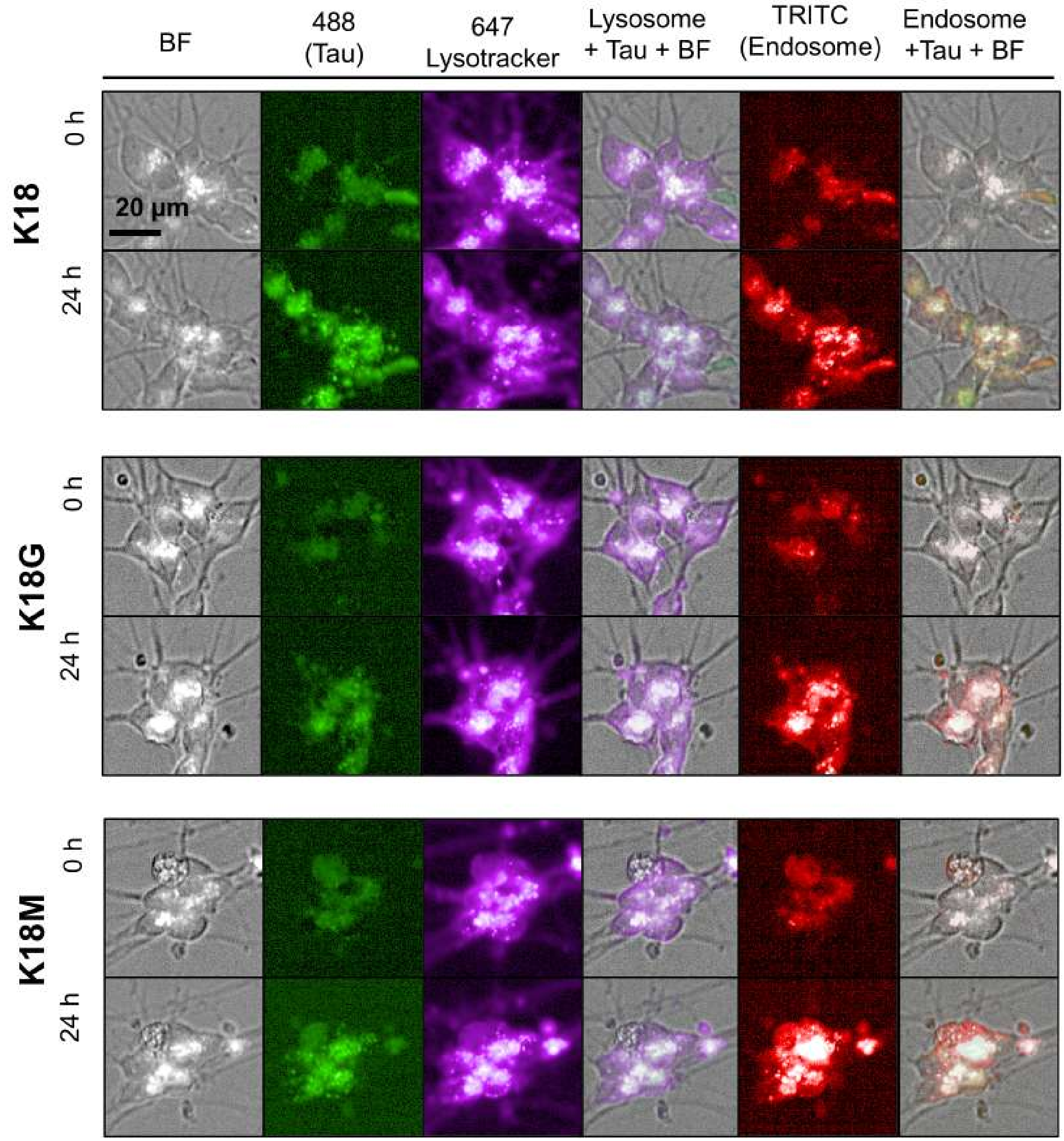

### 11. Colocalization of Aggregated K18 Tau with Early Endosome Tracker

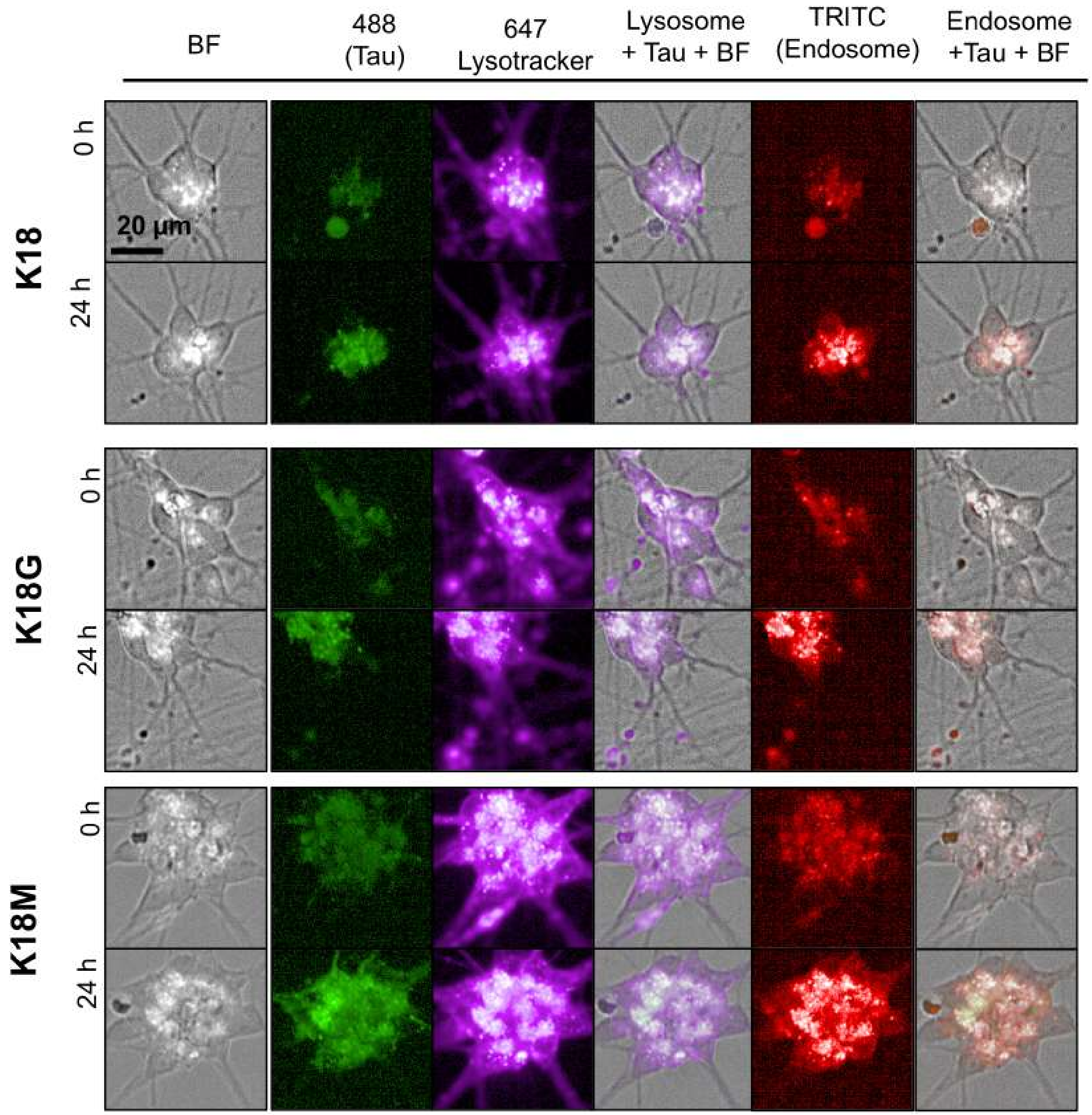

### 12. Colocalization of Monomeric 2N4R Tau with Late Endosome Tracker

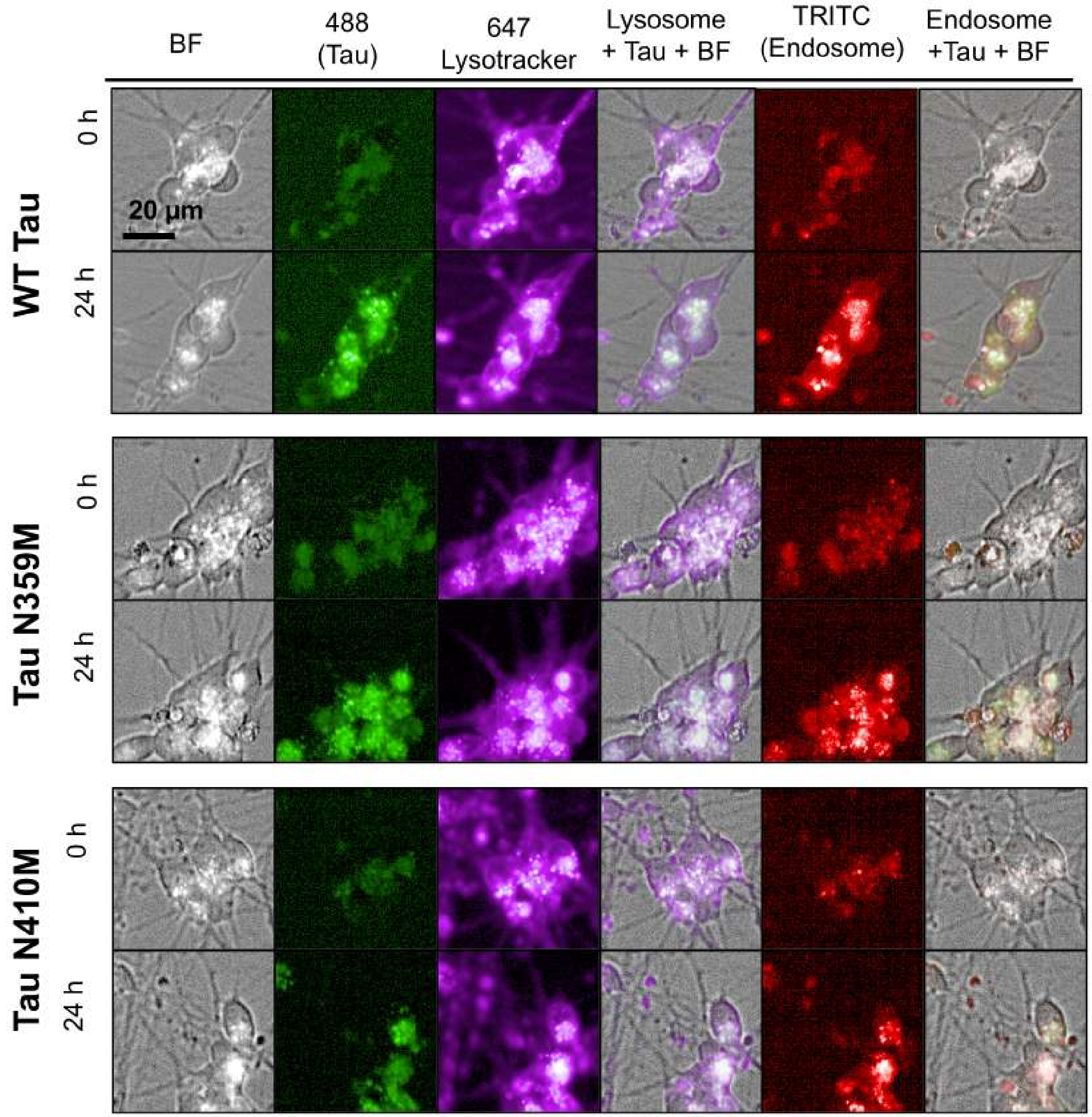

### 13. Colocalization of Aggregated 2N4 Tau with Late Endosome Tracker

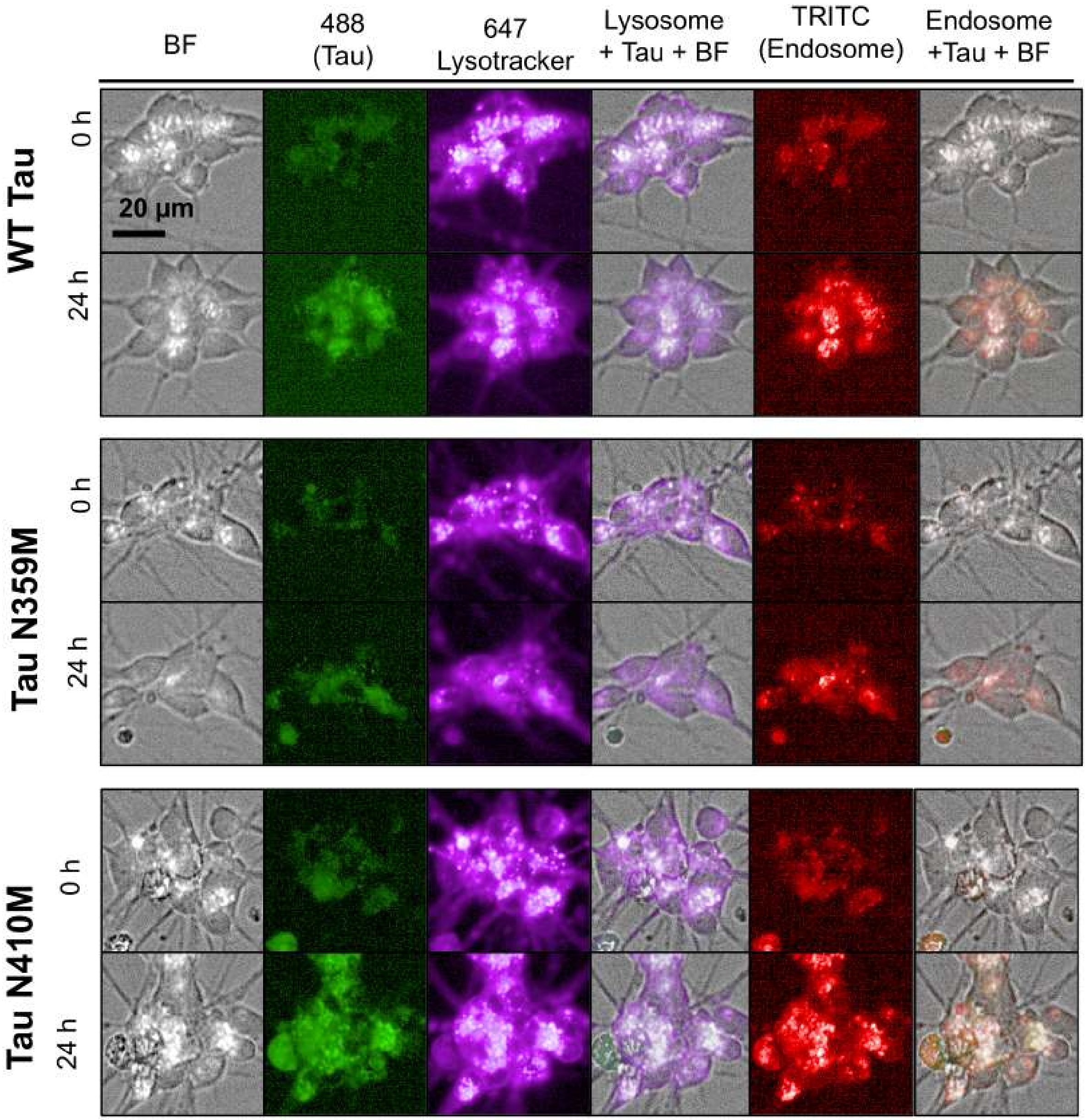

### 14. Colocalization of Monomeric K18 Tau with Late Endosome Tracker

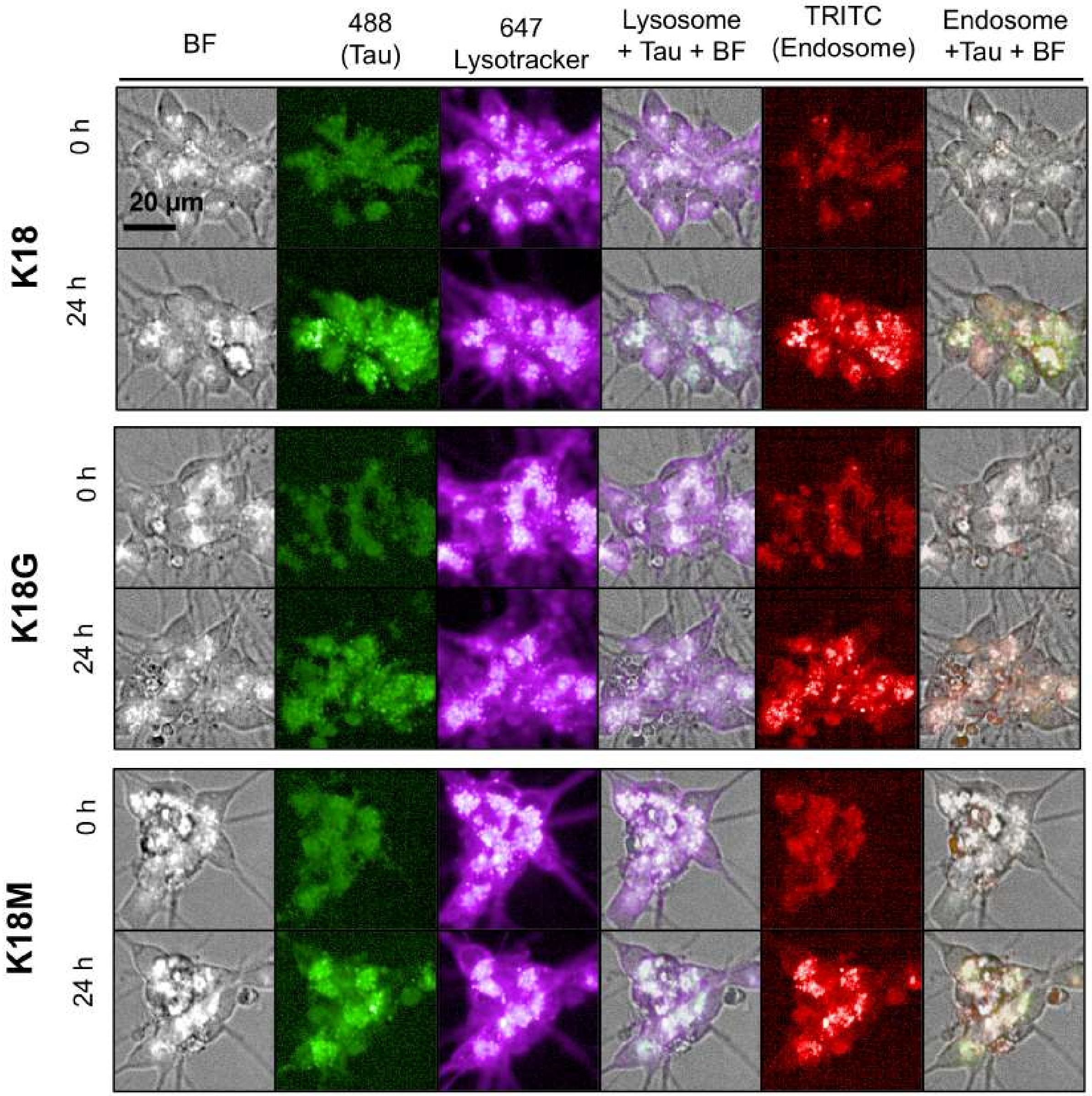

### 15. Colocalization of Aggregated K18 Tau with Late Endosome Tracker

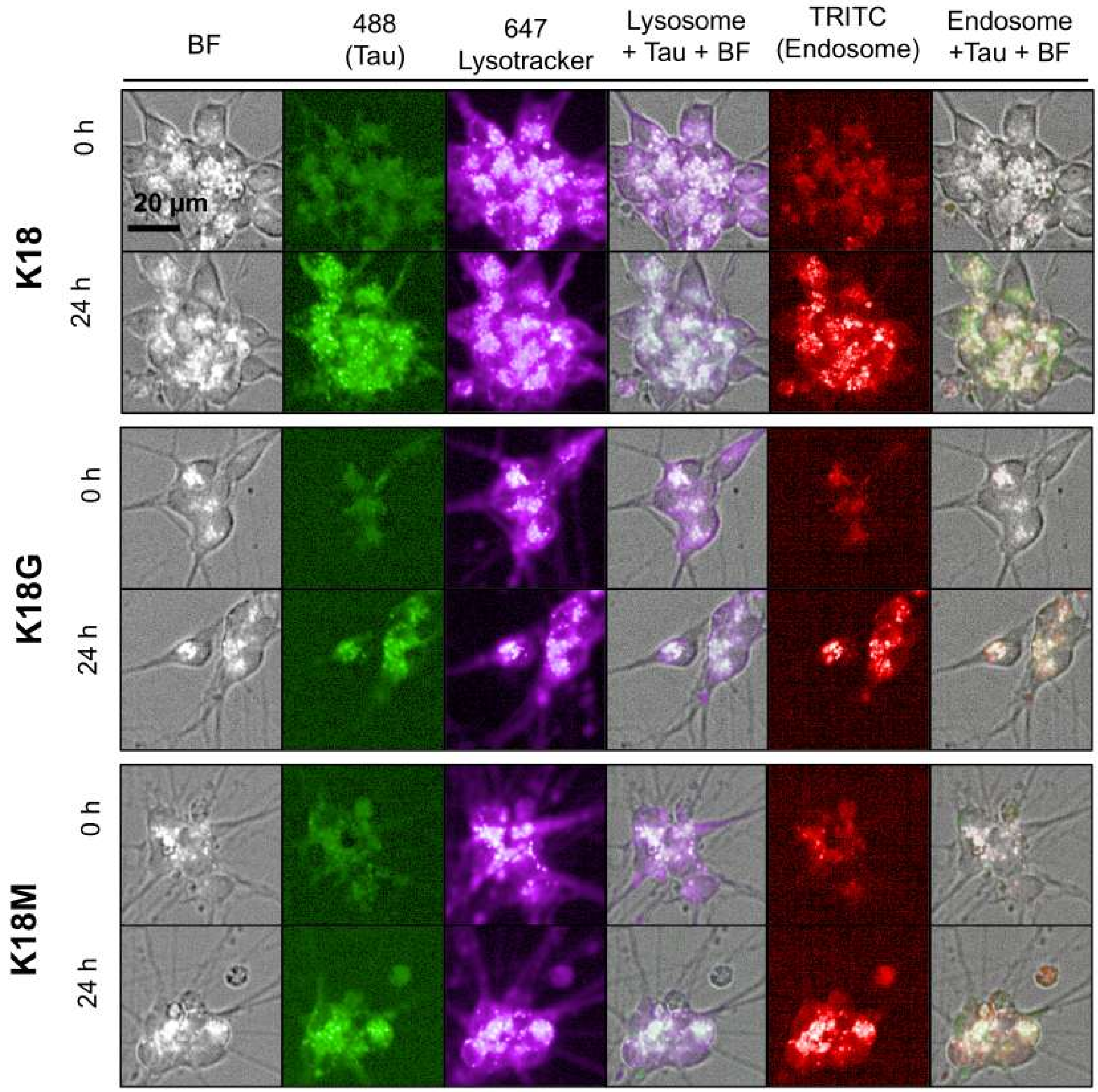

Experimental details: CellLight Early Endosomes-RFP (Invitrogen #C10587) at finial concentration 50 nM, and CellLight Late Endosomes-RFP (Invitrogen #C10589) at finial concentration 50 nM. LysoTracker Deep Red (Invitrogen #L12492) at finial concentration 50nM or LysoTracker Green DND-26 (Invitrogen #L7526) at finial concentration 50 nM.
